# Scaling up to meet new challenges of brain and behaviour through deep brain electrophysiology in freely moving sheep

**DOI:** 10.1101/2021.08.04.454778

**Authors:** Nicholas Perentos, Marino Krstulovic, A Jennifer Morton

**Author notes:** Professor A.J. Morton, Department of Physiology, Development and Neuroscience University of Cambridge, Downing Street Cambridge CB2 3DY.

## Abstract

While rodents are arguably the easiest animals to use for studying brain function, relying on them as model species for translational research comes with its own sets of limitations. Here, we propose sheep as a practical large animal species for *in vivo* brain function studies performed in naturalistic settings. To demonstrate their experimental usefulness, we performed proof-of-principle deep brain electrophysiological recording experiments from unrestrained sheep. Recordings were made from cortex and hippocampus both whilst sheep performed goal-directed behaviours (two-choice discrimination tasks), and across states of vigilance that included natural sleep. Hippocampal and cortical oscillatory rhythms were consistent with those seen in rodents and non-human primates, and included cortical alpha oscillations during immobility, hippocampal theta oscillations (5-6Hz) during locomotion and hippocampal sharp wave ripple oscillations (∼150 Hz) during immobility. Moreover, we found clear examples of neurons whose activity was modulated by task, speed of locomotion, spatial position, reward and vigilance states. Recordings were conducted over a period of many months. Due to the exceptional stability of individual electrodes we were able to record from some neurons repeatedly across the span of a month. Together these experiments demonstrate that sheep are an excellent experimental animal model to use in longitudinal electrophysiological and imaging studies, particularly those requiring a large brained mammal, large scale recordings across distributed neuronal networks, experimentation outside the confounds of the traditional laboratory, or all the above concomitantly.

## Introduction

Electrophysiological studies conducted *in vivo* using rodents have opened up new vistas in our understanding of brain function during behaviour. By using suites of advanced neural recording techniques (Ghosh et al., 2011; Jun et al., 2017) and novel perturbation technologies (Kim et al., 2017) combined with theoretical and computational advancements that rely on large scale data (Paninski and Cunningham, 2018), our understanding of how different brain regions contribute to behaviour has never been greater. Intriguing as they are, however, mice are small nocturnal mammals that bear little resemblance to humans in either brain structure (Dawson et al., 2018) or behaviour. Furthermore, there is growing concern about the limitations of rodents as pre-clinical translational models, since therapeutic interventions that are successful in mice can perform poorly in human clinical trials (Atkins et al., 2020; Dawson et al., 2018; Drummond and Wisniewski, 2017; Leenaars et al., 2019; Nestler and Hyman, 2010). Indeed, the arguments in favour of diversifying the species used in neuroscience research are numerous. Apart from translational reasons, that may be particularly pertinent for study of brain dysfunction (Hodge et al., 2019), limiting investigations to a small set of species leaves the experimenter unaware of potential alternatives solutions that Nature offers to challenging biological problems (Keifer and Summers, 2016; Yartsev, 2017). Recent advances in gene editing technologies, combined with the development of neural interfaces and imaging technologies that have cross species applicability, mean that this is an ideal time to explore the potential for using larger mammals as model organisms (Eaton and Wishart, 2017; Fan et al., 2018; Pouladi et al., 2013).

To bridge the gap between rodent and non-human primate animal models, we have been investigating the use of sheep as a model species for functional brain research. In the behavioural domain, despite their reputation for limited cognitive ability, sheep perform better than mice in many forms of cognitive testing (McBride et al., 2016; Nicol et al., 2016; Sorby-Adams et al., 2021) and have a high predicted level of cognitive capacity that approaches or equals that of non-human primates (McBride and Morton, 2018). At the gross anatomical level, particularly with respect to the basal ganglia, the sheep brain is much more similar to the human compared to that of rodents. Furthermore, their diurnal lifestyle and similar sleep physiology means that sheep sleep studies are directly relevant to humans (Morton et al., 2014; Morton and Howland, 2013; Schneider et al., 2021; Vas et al., 2021). To date, however, recordings at single neuron resolution from sheep have been conducted only under anaesthesia or during head-clamp-restrained conditions (Clarke et al., 1976; Gierthmuehlen et al., 2014; Kendrick and Baldwin, 1989). The potential for experimentation in large gregarious mammalian species that can not only be used safely when unrestrained but can also be easily trained to perform behavioural tasks arguably allows a unique opportunity to study brain function during complex behaviours such as learning and socialising. For these studies we adapted a well-characterised single unit recording method, using arrays of individually adjustable tetrodes allowing micrometre level resolution in electrode placement. This method is well established in several other species including rodents (e.g. (Nguyen et al., 2009) and non-human primates (Kyle et al., 2019), and allows long-term recordings from virtually any brain structure.

We describe for the first time in sheep several electrophysiological signatures at the level of both network dynamics, via local field potentials, and single neurons. These include oscillatory network dynamics in cortex (e.g. alpha band oscillations) and hippocampus (e.g. theta and sharp wave ripple oscillations) that are largely similar to those observed in other mammalian species. As well we observed single neurons that are modulated by performance in cognitive tasks. The possibility of making stable recordings from single neurons over long periods creates new opportunities for studying brain function in naturalistic settings.

## Results

Recordings were made using tetrode microdrives (Fig.1a-d), with 16 and 24 tetrodes (in sheep 1 and 2, respectively) implanted onto the dorsal aspect of the cranium above the ectolateral gyrus (For details, see Methods, Suppl. Fig. 1). During testing, sheep were tethered to the recording system by a cable that allowed unimpeded movement within the experimental arena (Fig. 1e-h, Suppl. Video 1-4). Typical recording sessions started with a baseline recording phase during which the sheep was restrained outside the behavioural arena. Following this baseline period was a phase of active behaviour that would typically take place inside the experimental arena. This phase comprised cognitive testing (Fig. 1e, Suppl. Video 1,2) or exploration/foraging (Fig. 1f).

**Figure 1.**
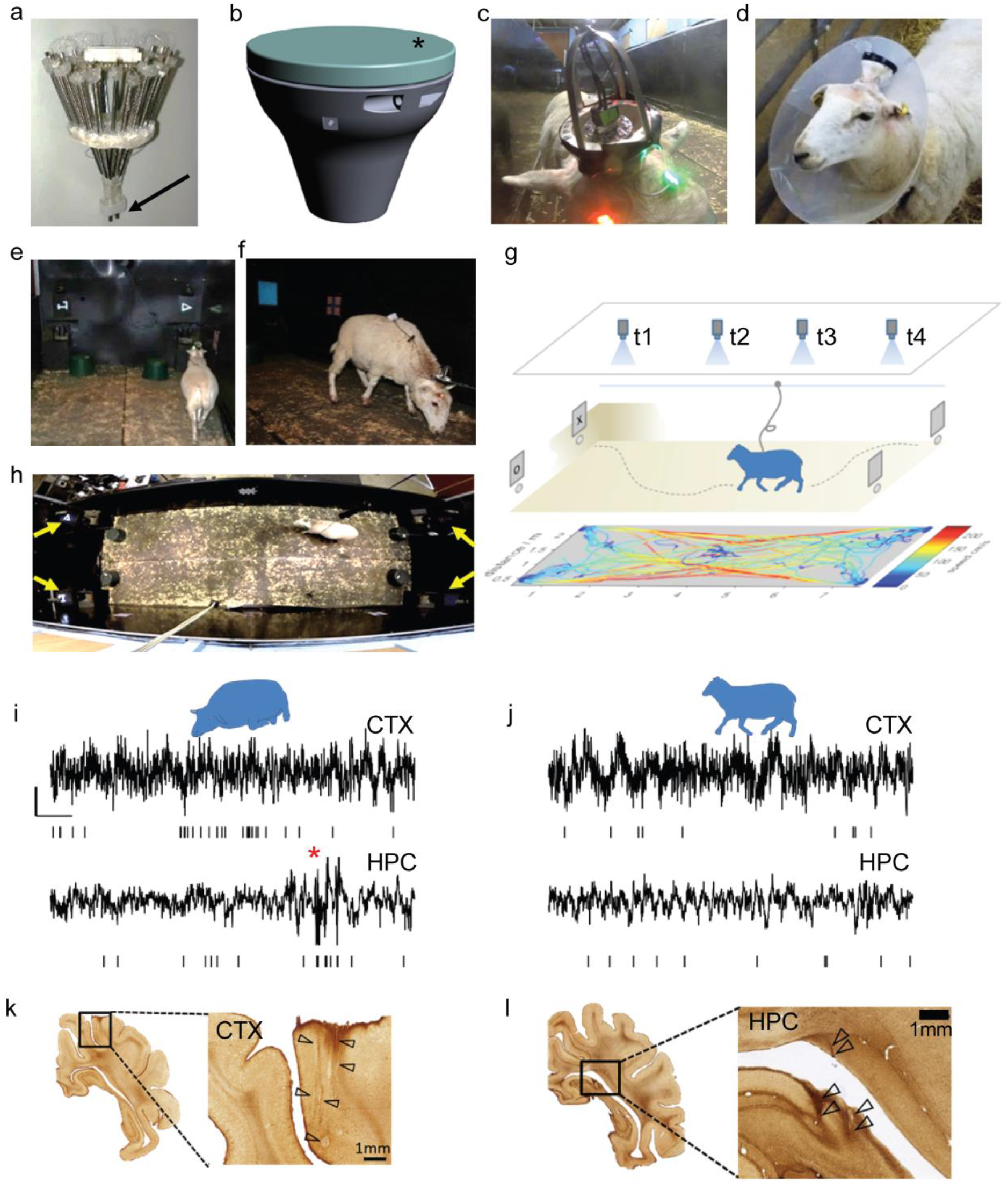
Apparatus and experimental procedures for single neuron and local field potential recordings in freely moving sheep. (a) Custom-made 22-tetrode microdrive assembly before implantation shows tetrodes terminating in two cannulae (black arrow). The microdrive is housed in a 3D-printed chamber (b) that is secured onto the skull. (c) During recording, the cap of the protective casing (* in b) is removed, and a metal frame that anchors the recording cable and protects the connection is attached. During post-operative care and recovery, the implant and surgical incision is further protected using an Elizabethan collar (d). During recording, sheep either perform in a two-choice discrimination task (e), or forage for food pellets (f) dispersed on the floor of the arena. The schematic (g) shows the position of overhead cameras (t1-t4) that are used to record the position of the sheep inside the arena and an example positional tracking trace that is color-coded by the speed of locomotion (colour scale corresponds to 0–200 cm/s). (h) An aerial view of the experimental arena showing the position of the four computer monitors used for two-choice discrimination testing (yellow arrows). Example local field potential traces shown were recorded from cortex and hippocampus during restraint (i) and locomotion (j). Vertical ticks below each trace represent putative spike events. The red star in (i) indicates a sharp wave ripple event. Example histological sections used to verify electrode positions at the end of experiments in cortex (k) and hippocampus (l). CTX = cortex, HPC= hippocampus.

During each recording session there was also a post-testing phase during which the sheep was restrained for an additional recording of immobility related brain activity as well as electrode advancement. Fig. 1i and j show examples of local field potential (LFP) and multiunit spiking activity recorded from tetrodes located in the cortex or the hippocampal formation. During restraint/immobility (Fig. 1i), cortical activity exhibited a rhythmic LFP, whereas the hippocampal activation was dominated by slow and irregular activity. As expected, during locomotion and active behaviour, the cortex exhibited less rhythmicity in comparison to the immobility intervals, while the hippocampus exhibited prominent oscillations. These observations are quantified in detail below. At the end of all experiments the final location of a subset of tetrodes was marked with electrolytic lesions and electrode locations were verified through post-mortem histological examinations (Fig. 1k and l).

### Stable extracellular recordings are feasible in freely moving sheep

We recorded from two sheep daily over a period of several months (∼8 and ∼6 months from sheep 1 and 2, respectively) during which time we identified a total of 3308 putative single neurons (Fig 2). Each day we were typically able to record from several neurons at a time (see clusters from an example tetrode (Fig 2a), the associated waveforms (Fig 2b and example autocorellograms in Fig 2c). The maximum number of identified putative single neurons (henceforth referred to as neurons) on a single day was 36 while the average for sheep 1 was 10.6 ±0.5 and for sheep 2 was 16.5±0.7. As assessed by the L-ratio and isolation distance metrics (Fig. 2d), the quality of cell recordings reflected those previously reported in the literature (Schmitzer-Torbert et al., 2005). Given the absence of any intracellular recordings from sheep neurons with which to make a comparison, we opted to subdivide putative neurons into the two broad categories based on the duration of the after hyperpolarisation (dAHP) as has been done previously with recordings from non-human primates (Fan et al., 2017) rodents (Sirota et al., 2008) or guinea pigs (McCormick et al., 1985). We thus classified cells into narrow (dAHP < 225 (μs) and broad (dAHP > 250 (μs) spiking neurons (Fig. 2e,f). Cells within the transition zone between these two categories were treated separately (see below). A small subset of neurons that appeared highly symmetrical were classified as potential ‘axonal’ events (Fig. 2f, inset). Average firing rates were predominantly low with most cells (∼80%) exhibiting rates at or below 5 Hz (Fig 2g) while firing rates based on the above categorisation revealed a statistically significant higher probability of higher firing rates within the narrow spiking category in comparison with the broad spiking category (Fig. 2h). This latter observation is consistent with a categorisation that could be capturing a principal pyramidal vs interneuronal division (Fan et al., 2017; McCormick et al., 1985; Sirota et al., 2003).

**Figure 2.**
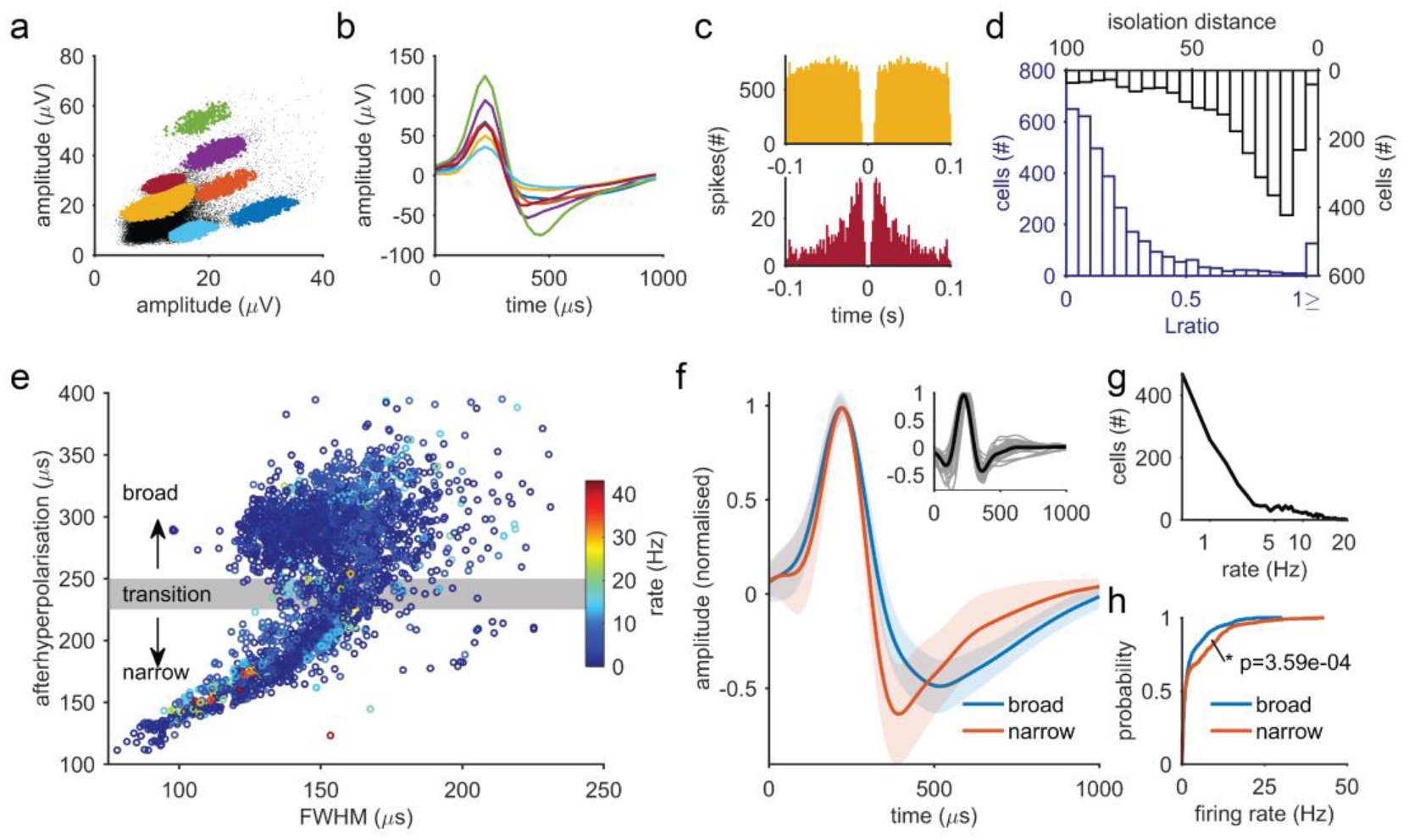
Characteristics of extracellular recordings of neurons from sheep cortex and hippocampus. (a) An example recording from a tetrode showing seven identified single unit clusters. (b) Average spike waveforms for each cluster from the recording shown in (a). (c) Example autocorrelograms from two single units from the recording shown in (a). (d) Distributions of cluster quality characteristics (isolation distance and L ratio) using recordings from both sheep. (e) Graph showing the entire population of recorded neurons plotted according to their extracellular waveform shape characteristics. Waveforms were grouped into narrow (<225μs) and broad (>250μs) on the basis of the delay from the waveform maximum to the most negative peak after the maximum and shown in (f). The solid orange and blue lines in (f) show the mean spike waveform of the narrow and broad populations respectively (light orange and blue are the corresponding standard errors of the means). Inset in (f) shows the mean waveform of a small subset of cells n = 32 with symmetric waveforms. (g) Population-wide distribution of firing rates. (h) Cumulative distribution functions for broad and narrow waveforms. FWHM = full width at half maximum.

With time, a decreasing number of neuron yield was observed. This was likely due to a combination of technical factors that include the location of electrodes (since, for example, later recording days included tetrodes that were travelling through white matter or were located in the ventricles), deterioration of electrode impedance, and physical damage leading to the loss of tetrodes connections. Nevertheless, even after several months, we obtained good quality recordings. At later stages in the study, where stable clusters of high signal-to-noise ratio were identified we attempted to record from these groups across multiple days by not advancing the tetrodes. In such cases we were able to record from clusters whose firing rate, peristimulus time histograms and autocorrelograms were stable across days. The longest period of continuous recording from identified clusters was 48 days (Suppl. Fig. 2).

### Cortical single unit correlates of behaviour during two-choice discrimination

To explore the neural correlates of behaviour within the experimental arena, we used a two-choice discrimination behavioural paradigm in which an automatic testing apparatus was used (Fig. 1e,h). The sheep was required to approach the apparatus at one end of the arena and choose one of two automatically displayed stimuli, in order to obtain a pellet feed reward that was dispensed upon the correct choice (S+). Approach to the alternative stimulus (S-) was not rewarded and was additionally followed by a high-pitch auditory cue. An identical two-choice discrimination apparatus was installed at the opposite end of the experimental arena. These were activated alternatively, so the sheep was required to traverse the arena in order to reach the next reward opportunity. We used cortical recordings made during these experiments to explore possible neural correlates with the behaviour., Performance with respect to the behavioural task was consistent with previous work in sheep (McBride et al., 2016). Even though sheep were neither food- nor water-deprived, they worked willingly and were able to learn new symbol associations daily (Fig. 3a, Suppl. Fig. 3).

**Figure 3.**
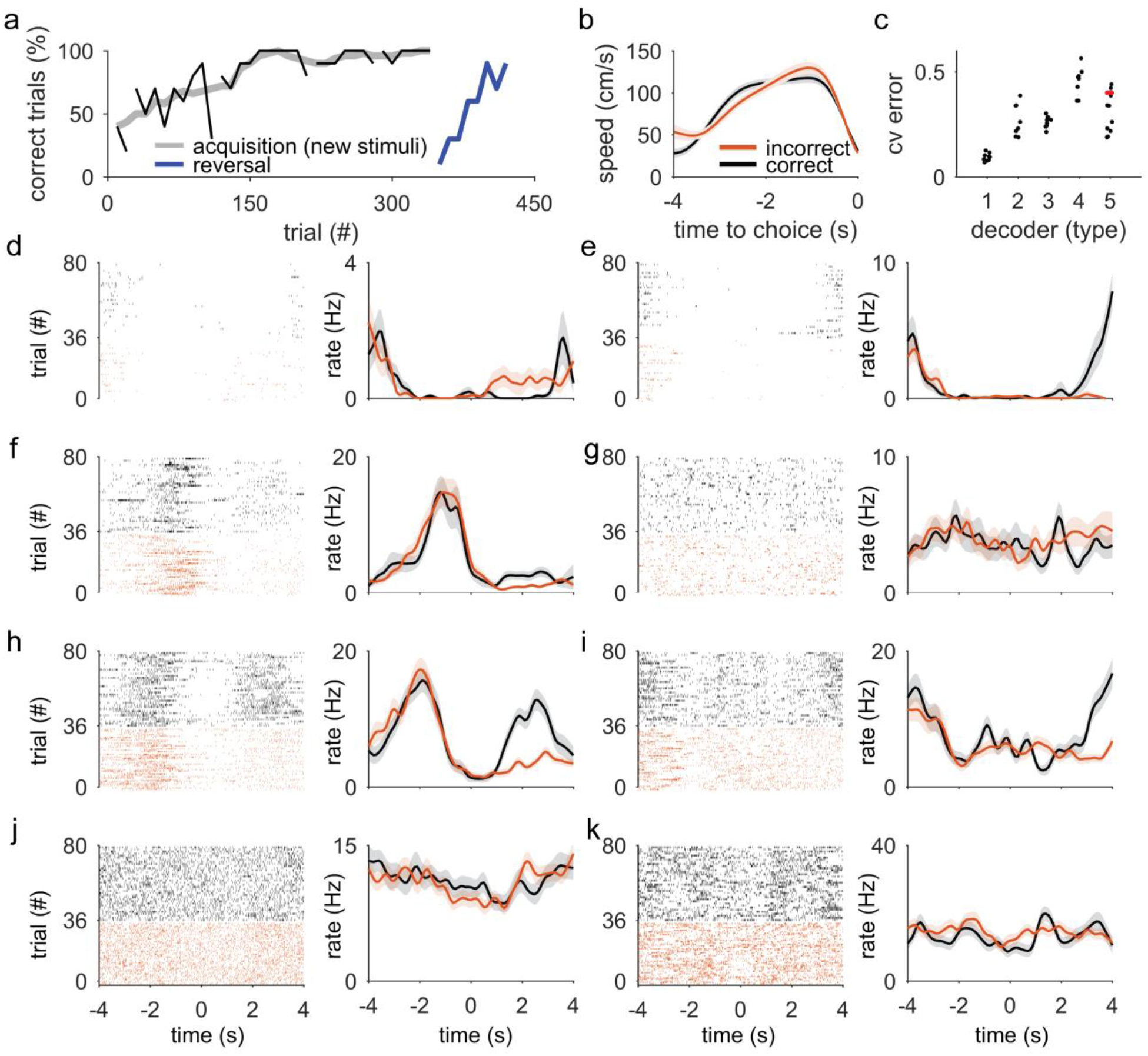
**Single neuron correlates of behaviour during two-choice discrimination**. (a) Serial two-choice discrimination performance across 6 consecutive days. Days 1-5 correspond to an acquisition stage (black lines and grey line moving average across days 1-5) and day 6 (blue) corresponds to performance at reversal of contingencies. (b) Speed profiles on approach to the choice points for correct (black) and incorrect (orange) trials. (c) Decoding accuracy for several behavioural variables from population firing rates corresponding to time windows preceding choice time points. Decoder types are 1: locomotion vs immobility, 2: task type (two-choice discrimination versus stimulus association, 3: speed of locomotion, 4: location of current choice point (1 of 4 choice points) and 5: correct versus incorrect choice). The datapoint in red corresponds to the session for which the raster plots are plotted in (d-k). (d-k) Raster plots from 8 example cells (left) and corresponding trial-averaged mean firing rates (right) centred onto the choice time-points (t =0s). Lighter grey and orange shades in firing rate graphs represent the standard error of the mean. Trials are grouped by choice outcome (black rasters= correct; orange rasters= incorrect). cv error: mean cross validation error.

We selected a subset of all recordings to analyse during the above described behaviour, choosing recordings that had several neurons simultaneously recorded as well as adequate positional tracking that allowed reliable estimation of the speed of movement. We asked which of the measured behavioural variables best predicted population activity of this cortical area. The behavioural variables were ‘task type’ (simple two choice discrimination with only one symbol displayed at a time, versus conventional two-choice discrimination, where the sheep had to choose between two symbols), ‘choice outcome’ (correct versus incorrect choice), ‘location’ (identity of the reward dispenser being accessed, i.e. spatial position at choice time-point), ‘speed of movement’ (shown in Fig. 3b), and ‘locomotion’ (whether the sheep was locomoting or not). As evidenced by the cross-validation error of each decoder (Fig. 3c) most recorders performed poorly except for the one for locomotion versus stationary conditions. This is perhaps unsurprising given the differential state the sheep is expected to be in during these two conditions (locomoting versus consuming reward, for example). The observation would also be consistent with results from recordings in the primary visual areas of the rodent cortex where behavioural modulation by speed of locomotion and other movements clearly influence activity in such primary sensory areas (Niell and Stryker, 2010; Stringer et al., 2019).

Simultaneously recorded cortical cells varied in their firing rates around choice points, with some cells showing increased and others decreased firing rates following choice. Eight examples of simultaneously recorded cells recorded during two-choice task performance are shown as raster plots zeroed around the decision point (Fig. 3 d-k). While the firing rates of each cell was characteristic on approach to correct and incorrect trials, following the choice the firing rate some of them diverged.

### Neuronal signatures of quiet wakefulness and sleep in sheep

During immobility ’baseline’ recordings, the sheep behaved in a relaxed manner with limited movements and no struggling. Their eyes were sometimes closed but mostly open. During this phase, electrophysiological features consistent with quiescence/inactivity were observed in both hippocampus and cortex (Fig.4a-l). For example, sharp wave ripple (SWR) oscillations of the hippocampus are high frequency oscillatory events (>100Hz) that have been observed in many mammalian species and are associated with memory consolidation processes (Buzsáki, 2015).We confirm here for the first time that SWRs are also a prominent feature of the sheep hippocampal formation during immobility (Fig. 4a-g). Both components of this network oscillation formed by the slower sharp wave component together with a high frequency oscillation of ∼150Hz were readily observable across ∼8000 events and multiple recording days (Fig. 4b,c). In the hippocampus, during SWRs there was a statistically significant higher probability of spiking at ripple onset (Wilcoxon signed rank test, Zmin >3.62, P<0.002; Fig. 4d, Suppl. Fig. 5). In contrast, no such modulation was observed in cortical spiking activity (Wilcoxon signed rank test, Zmax >0.97, P>0.2; Fig. 4e). Mean SWR duration was ∼70ms while the mean frequency of occurrence was 15 events/min (Fig. 4f,g).

**Figure 4.**
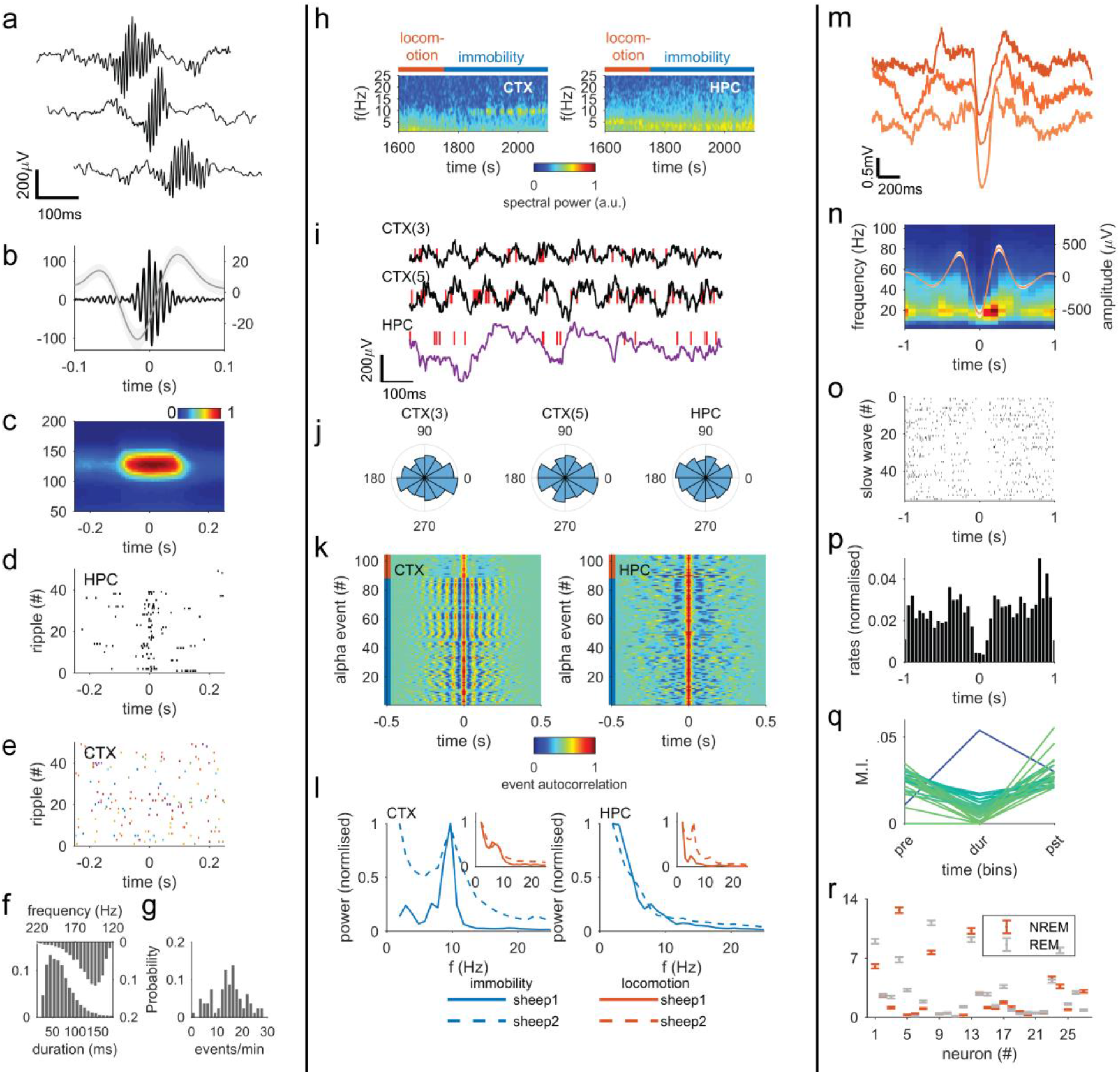
Electrophysiological profiles of sheep cortex and hippocampus during immobility and sleep. (**a-g**) Hippocampal (HPC) sharp wave ripple (SWR) oscillations are observed during immobility. (a) Example SWR oscillations. (b) Average sharp wave (grey curve, shaded region represents the SEM) component and the average ripple component (black) of the SWR. (c) Ripple-triggered average spectrogram of all ripples detected in sheep 1. A prominent spectral peak is observed at ∼140 Hz. SWR-peak triggered raster plots of single unit activity in the HPC (d) and CTX (e). (f) Top: distribution of peak repetition rates of ripple oscillation (µ = 155.1 Hz, µ_1/2_ = 150.7 Hz), bottom distribution of ripple duration (μ = 69.1 ms, µ_1/2_ = 63.5 ms). (g) Distribution of SWR frequency of occurrence (µ = 14.6 events/min, µ_1/2_ = 15.0 events/min). (h-l) Cortical alpha-band oscillations. (h) Example spectrograms from the same sheep and recording from cortex (left) hippocampus (right). (i) CTX and HPC LFP during detected alpha-band oscillation events. Red ticks represent detected spikes. (j) Polar histograms of three example units’ activities with respect to the phase of alpha oscillations. (k) Detected alpha band oscillations in the cortex (left) and their corresponding activities in the hippocampus (right). (l) Normalised spectral power for the states of immobility and locomotion in CTX (left) and HPC (right). (m-r) Neural dynamics during sleep. (m) Examples of detected slow wave oscillations. (n) CTX spectrogram triggered onto detected slow-waves. Superimposed is the average slow wave waveform (shaded area around average is SEM). (o) Example raster plot of a cell exhibiting clear down state around 0s. (p) Mean activity across all cells around detected slow-wave events. (q) Mean firing rates of cells before, during and after slow-wave events. (r) Mean firing rates for all cells during REM and NREM sleep. CTX = cortex, HPC= hippocampus.

Next, we focused on cortical activity during quiet (immobile) behaviour. We observed prominent ∼10 Hz oscillations that were absent during periods of active behaviour (Fig. 4h). These oscillations modulated local spiking activity and occurred in synchronisation with large irregular activity in the hippocampus (Fig. 4i-k). Both sheep exhibited similar power spectra for both locomotion and immobility/quiescence periods (Fig. 4l and Suppl. Fig. 4). Finally, to identify electrophysiological signatures of sleep, we obtained an overnight recording from sheep 2 during which we observed prominent slow oscillations within the cortex (Fig. 4m). These events occurred while the sheep showed all signs of having fallen asleep (being recumbent, immobile and with eyes closed). To corroborate whether these events were *bona fide* local slow wave oscillations of non-rapid eye movement (NREM) sleep, we searched for two known correlates of these sleep events. First, we searched for spindle-like events that would be phase locked to the slow oscillation (Fig. 4n). We found elevated ∼20 Hz activity at the transition between slow wave negativity-to-positivity was observed that is consistent with thalamocortical spindles as observed in many other species (Siegel, 2005). Second, we examined modulation of spiking activity within a 2-second window around the peak negativity of slow waves. This revealed a statistically significant reduction in multiunit spiking activity near the peak negativity (down-state; Wilcoxon signed rank test, Zmax <-3.91, P<0.001; Fig. 4o-q). Finally, we observed periods of sleep that were consistent with rapid eye movement (REM) sleep that was characterised by both increased and decreased firing rates of single units (Fig. 4r, see also Suppl. Fig. 6).

### Hippocampal dynamics in freely-moving sheep

An apparently ubiquitous property of the hippocampus during alert states across species is the 4-10 Hz theta oscillation (Buzsáki et al., 2013). We were able to show for the first time that such rhythms are readily observed in the sheep hippocampus together with gamma band oscillations (Fig. 5a) especially during periods of activity (Fig. 5a-c, Suppl. Video 3,4). Speed of locomotion is a robust behavioural correlate of theta oscillations in rodents. We therefore explored whether a similar correlation was present under our experimental conditions (Fig. 5 and Suppl. Figs. 6,7). As can be seen in Fig. 5e the distribution of speed during the two-choice discrimination task was bimodal with fewer counts for speeds between the two peaks. For further analysis we therefore partitioned speed data into three bins (low, <12 cm/s; intermediate,12 – 80 cm/s); high, >80 cm/s). We computed corresponding average spectra (Fig. 5f) followed by a statistically significant one-way ANOVA on the peak power within the 4-7 Hz range (F(2, 2443) = [4.361], p = 0.0129) with theta peak power significantly higher in high speed versus low speed, (p = 0.0103, 95% C.I. = [6.85, 68.51]) while no statistically significant differences were observed between low speed versus intermediate speed (p=0.118) nor between transition and high speeds (p=0.745). Regression analysis of theta power revealed only weak correlations with speed or acceleration (R^2^ = 0.22, p=2.54·10^-60^ and R^2^ = 0.24, P = 1.29·10^-144^, respectively). A stronger effect was observed for the peak power within the 10-13 Hz band corresponding to the first theta harmonic (one-way ANOVA F(2,2443) = 117.40, p = 2.07x10^-49^, with all three speed bins being statistically significantly different from each other (low vs transition: p < 0.01, 95% C.I. = [7.15,15.60], low vs high: p < 0.01, 95% C.I. = [23.80,32.86], intermediate vs high: p < 0.01, 95% C.I. = [13.02,20.89]). Similarly, gamma band activity increased progressively as a function of speed (Fig. 5g, one-way ANOVA F(2,1067) = 222.99, p = 1.21x10^-81^, with low vs high and intermediate versus high being statistically significant (low vs high: p < 0.01, 95% C.I. = [113.23,166.62], intermediate vs high: p < 0.01, 95% C.I. = [121.86,157.77] and low versus intermediate p = 1).

**Figure 5.**
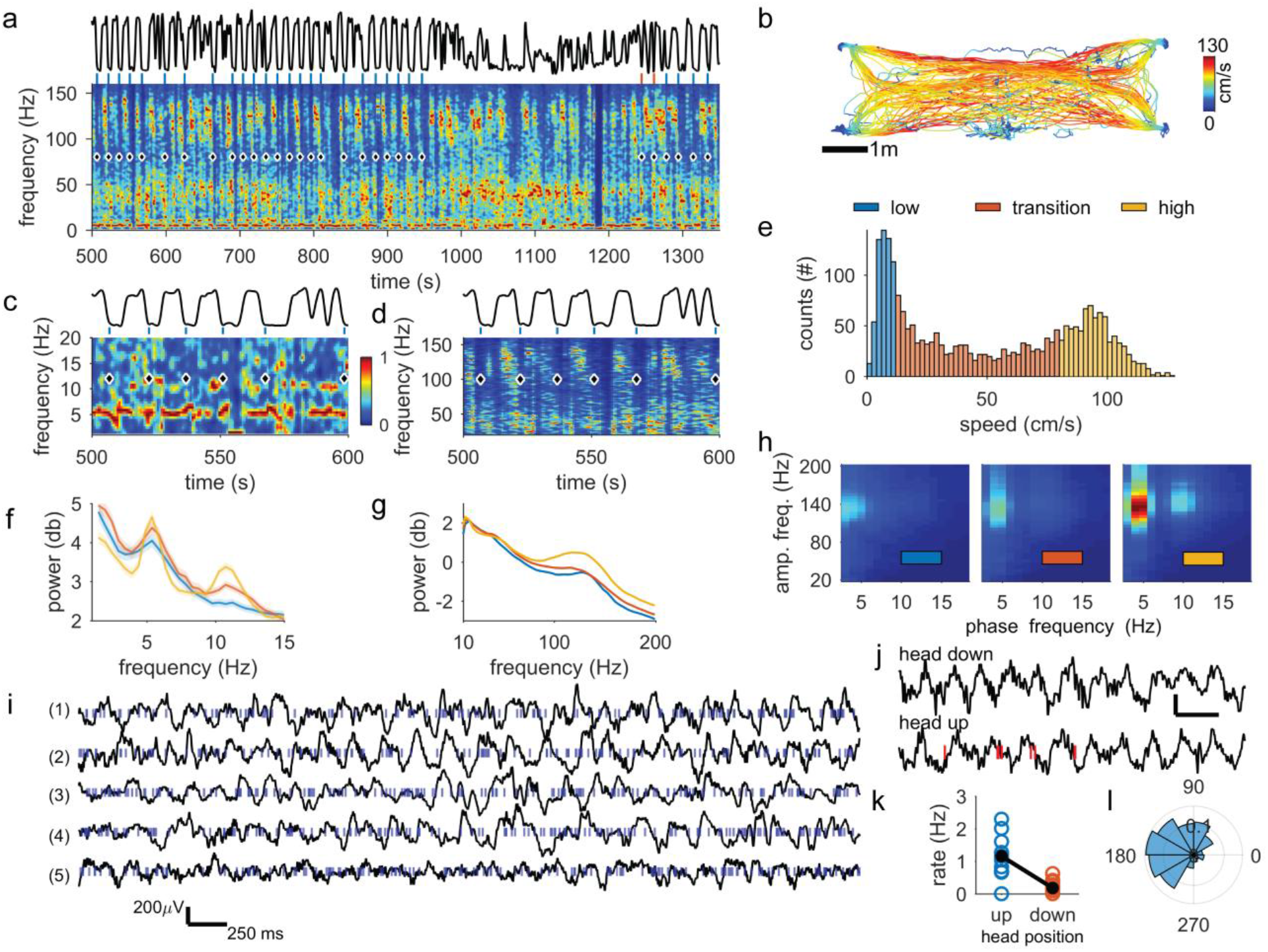
**Hippocampal oscillatory dynamics during cognitive testing**. (a) An example of a hippocampal (HPC) spectrogram during cognitive testing. Black trace at the top of the panel depicts the speed of movement and ticks above the spectrogram indicate choice time points (blue for correct and red for incorrect choices). (b) Speed profile is show as a function of position in the arena. Data correspond to the time interval depicted in (a). Zoomed in sections of (a) are shown in (c,d) for the 500 – 600s time period for theta (c) and gamma (d) bands. Black trace and ticks are as described in (a). (e) Histogram showing speed distribution partitioned into low (blue), intermediate (orange) and high (yellow) bands. (f) Average power spectra for each of the three speed bins for theta range frequencies. (g) Average power spectra for each of the three speed bins for gamma range frequencies. (h) Comodulograms for the three speed bins. (i) Example HPC LFPs (black lines) and multiunit activity (blue ticks) during various behaviours where large amplitude theta oscillations were observed including (1) rearing with front hooves onto arena wall; (2) gaze fixation onto experimenter; (3) reward consumption; (4) restraint outside the arena and (5) locomotion during cognitive task. (j) An LFP (black) and putative single neuron (red ticks) activity during a guided traversal of the area with head either facing upwards or downwards. (k) Firing rate of the neuron shown in (j) for head up (blue) and head down (red) events. (l) Theta phase modulation of spiking activity. All spectrograms and phase-amplitude plots have a normalised (0 to 1) pseudocolor scale as depicted in (c).

**Figure 6.**
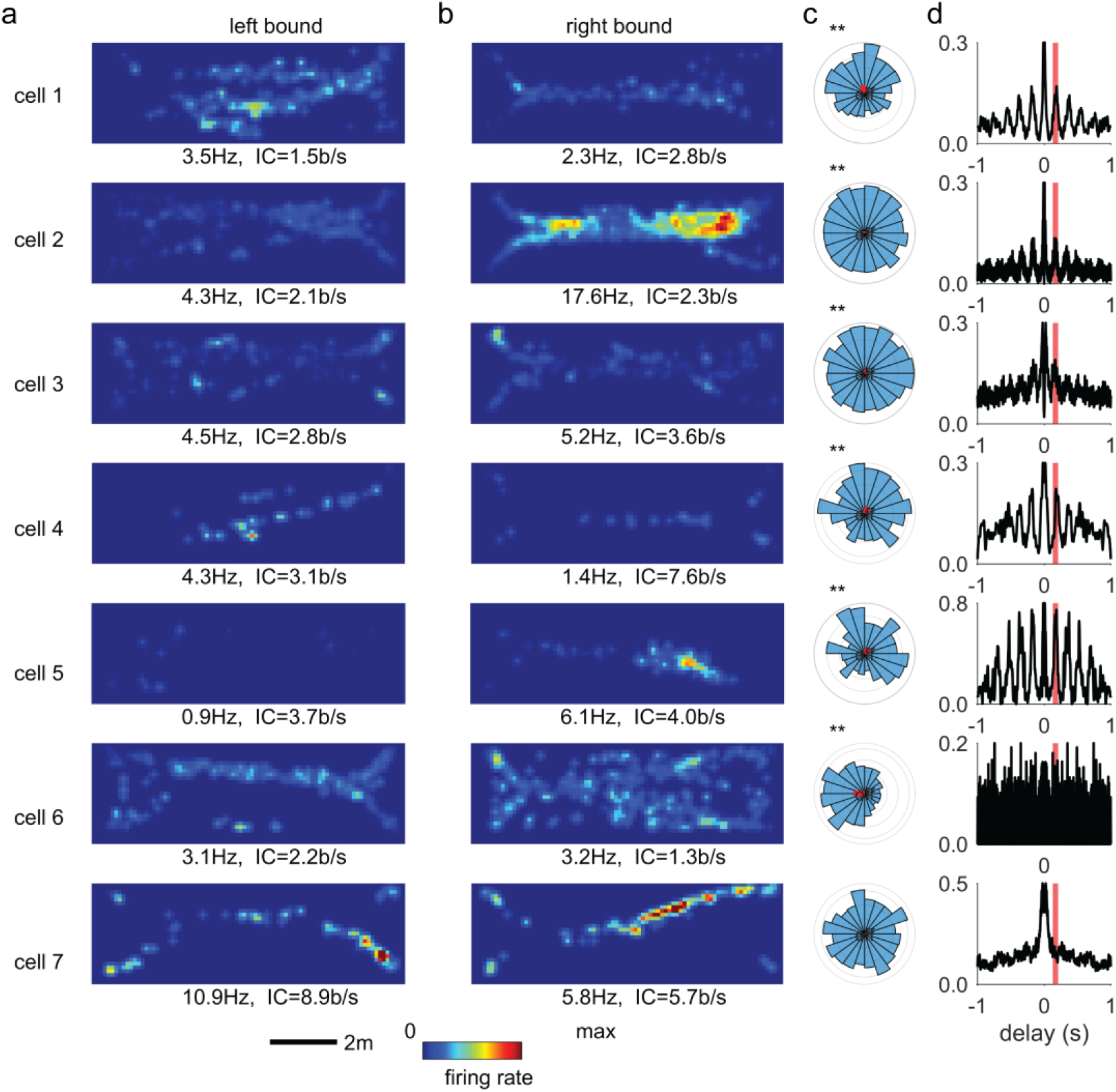
Hippocampal single unit activity in sheep is spatially modulated and entrained by theta oscillations. (a) Putative single neuron activity for left-bound trajectories. (b) Putative single neuron activity for right-bound trajectories. (c) Polar histogram of theta phase preference for each neuron. Resultant vector shown in red. (d) Autocorrelation function for each single neuron. Red line marks the delay that corresponds to theta frequency (170 ms). All cells except cell 7 show theta phase modulation. IC: Information Content. **: significant phase modulation at p<0.01.

**Figure 7.**
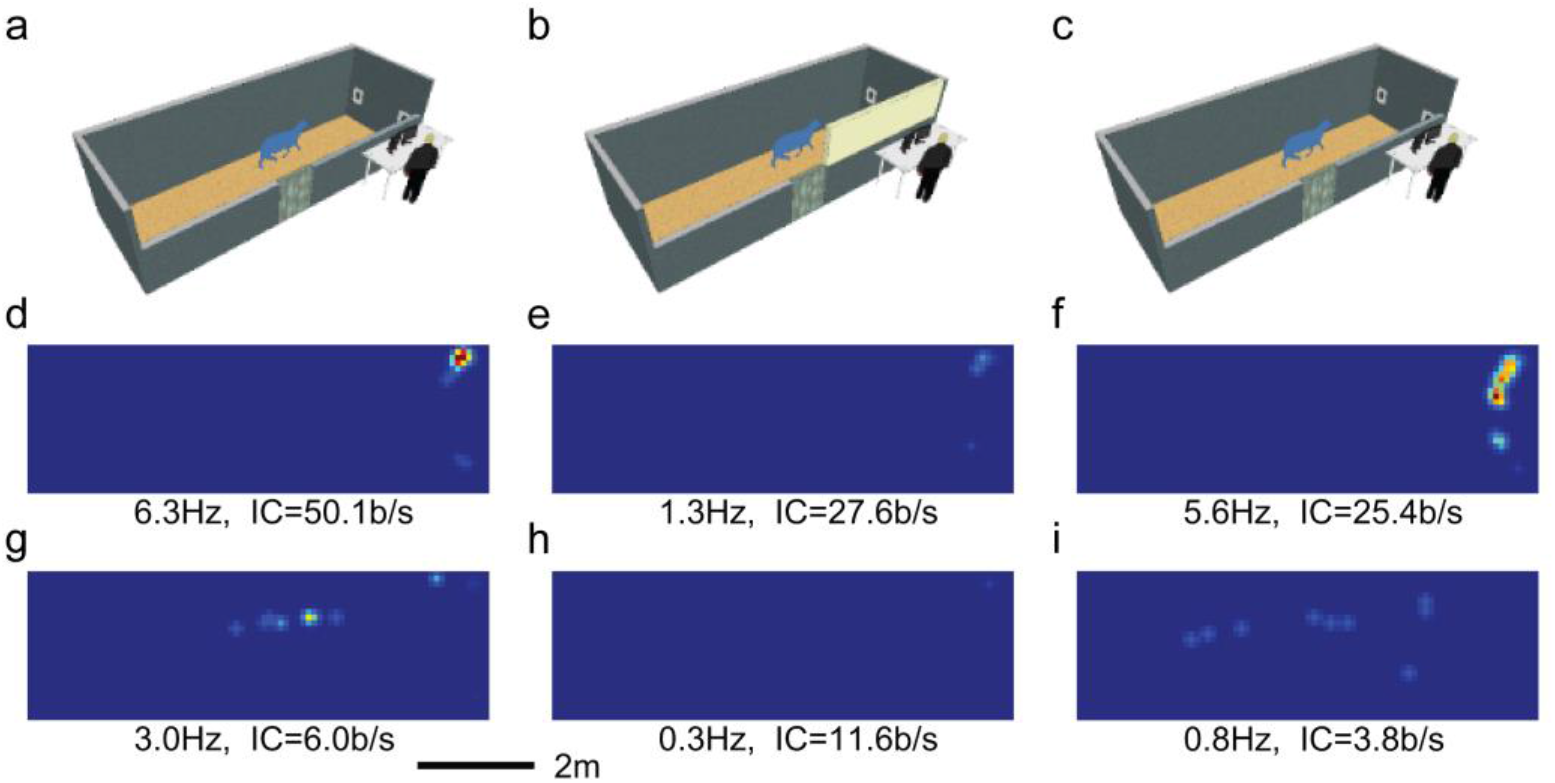
Rate remapping of a single neuron through distal cue manipulation. (**a-c**) Schematics show the experimental arena in different configurations. A wall extension (shown in **b** as a yellow rectangle) in the down (**a, c**) or raised (**b**) position forms the experimental manipulation. Note that the barrier was an extension of the arena wall made of the same material and color. (**d-f**) Spatial firing rate maps of a single neuron under different conditions within the same recording session. When the wall is down the neuron exhibits directional preference, with a higher firing rate for left-bound (upper heat maps **d, f**) than for right-bound (lower heat maps **g, i**) trajectories. When the barrier is raised to obscure the experimenter and other local cues (**b**), the firing rate of this cell is reduced significantly for both left- (**e**) and right-bound (**h**) trajectories. Once the barrier is lowered (**c**), firing rate for left-bound trajectories is restored to at least pre-manipulation levels (f). Each set of firing rate maps is constructed using a total of 40 approaches (20 left- and 20 right-bound). Firing rate and information metric for each cell is shown below each heat map. IC: information content.

Theta oscillations are widely reported to organise the power of gamma oscillations (Jensen and Colgin, 2007). Using a phase amplitude coupling quantification method (Tort et al., 2010) we observed that gamma band phase locking with theta oscillation phase increased as a function of speed bin (Fig. 5h). This effect was statistically significant for low versus high and intermediate versus high speeds (one-way ANOVA F(2,191) = 42.10, p = 7.10x10^-16^, low versus high: p<0.001, 95% C.I. = [0.135, 0.2446], intermediate versus high: p<0.001, 95% C.I. = [0.0986, 0.2082], and low versus intermediate: p = 0.4383) . Nevertheless, theta activity was readily observable in instances where the sheep was not locomoting, Fig. 5i. To visualise the behavioural correlates of theta further, we extracted periods in which theta oscillations were observed while the sheep was not locomoting. These typically occurred during occasional pauses in behavioural task, or whilst foraging for pellet rewards. Episodes associated with high power theta oscillations included rearing on hind legs, standing and staring at the experimenter, reward consumption and some occasions of restraint (Fig. 5i).

Finally, we noticed that some neuronal firing was modulated by differences in procedure (Fig. 5j-l). For example, when the sheep traversed the arena guided by the experimenter with its head pointing upwards, a theta- modulated neuron was preferentially active. In contrast, during guided traversal over the same path, but this time with the head facing forward/downward, this cell had a lower firing rate (Wilcoxon signed rank test, Z <3.33, P=8.7.10^-4^; Fig. 5j-l).

### Sheep exhibit spatially tuned hippocampal cells which can remap

In addition to the oscillatory dynamics in theta and gamma frequency bands already described, another hallmark of hippocampal activity is the spatial tuning of single cells (O’Keefe and Dostrovsky, 1971). We identified only a small number of putative hippocampal single neurons, because of electrode deterioration arising from the extended duration of recording. Nevertheless 7 of the 8 of the identified hippocampus cells we recorded were theta-modulated. Furthermore, 6 out of 7 of these cells showed spatial tuning with a directional preference (Fig. 6, cells 1-5, 7). That is, they appear to be ’place’ cells.

The activity of one additional hippocampal cell which was spatially modulated was further interrogated for possible remapping that could arise from manipulation of the spatial arrangements to the recording arena. During the behavioural testing and when the investigator entered the arena at the end of a testing period when the recordings were continuing, we noticed that this cell fired more when the sheep was looking at, or in the direction of, the investigator.

We wondered to what extent the experimenter served as a salient visual cue and whether their presence could modulate place cell activity. We therefore installed a barrier (an extension of the arena wall) that occluded the view of the experimenter to the sheep from within the arena (Fig. 7a-c). During a single recording, while the sheep was performing a cognitive task, we recorded the continuous activity of the place cell shown in Fig 7d-I (see also Suppl. Video 3). We recorded first with the barrier lowered and investigator visible (Fig. 7a), then with the barrier raised (obscuring the investigator; Fig. 7b) and then again with the barrier lowered (Fig. 7 c). We observed that the peak firing rate of this single unit dropped from 6.3 to 1.3 Hz when the barrier was up. After lowering of the barrier, the firing rate of this cell returned to levels similar to that seen in the first period of ’barrier-down’ recording. It therefore appears that this cell was sensitive to the visual cues available to the sheep. It remains to be explored whether this change is due to the rewarding nature of the experimenter’s presence or simply a visual sensory-related effect. It is an interesting possibility that this cell was responding to the ’social’ cue represented by the human experimenter.

## Discussion

In recent years, there has been a resurgence of interest in the use of large animals for the study of brain function. This stems from a growing awareness of increased translational potential of using larger animals such as pigs (Ulyanova et al., 2018) and sheep (Pfister et al., 2018) as preclinical models. In this study we show that it is possible to conduct deep brain recordings from sheep continuously over a period of several months. This wide timeframe would enable chronic studies both of pathology and therapeutic efficacy. Our data were collected from sheep that were unrestrained (apart from the tethering of the recording electrodes), either while they behaved normally (resting, locomoting, foraging, sleeping) or during the execution of laboratory behavioural tasks. We recorded simultaneously from CTX and HPC and characterised several thousand putative neurons, including some from the HPC that were modulated by place, and others that showed rate remapping after manipulation of sensory cues. The electrophysiological characteristics of neurons recorded this way were remarkably similar to those recorded from non-human primates. Our proof-of-principle study shows that sheep are a docile and tractable species in which to study brain function longitudinally. The ability of sheep to perform complex cognitive tasks (Sorby-Adams et al., 2021) coupled with new technologies that would enable multiple simultaneous recording sites, offers exciting new possibilities for understanding the mechanisms underlying only brain function in awake ’ behaving’ animals.

Historically, sheep have been an invaluable species for research into biological processes, particularly in the field of endocrinology and reproduction. With respect to neurophysiology of brain function, they have been less well used. We have used electroencephalography to study dynamics of sleep (Perentos et al., 2016a; Schneider et al., 2021, 2020) and the effect of neuroactive drugs (Nicol and Morton, 2020). At a deep brain level, notable examples of the use of sheep include the discoveries of orientation tuning or neurons in cortical areas V1 and V2 of adult sheep and lambs (Kennedy et al., 1983) as well as the description of facial recognition capacities and the underlying neural circuitry supportive of this function (Kendrick et al., 2001). These latter in vivo studies in sheep, however, were conducted in head-fixed or anaesthetised sheep. In this study, our aim was to demonstrate the potential of the species for neuroscience studies that require recording from individual neurons in awake and/or unrestrained ‘behaving’ sheep. As face validation for our techniques, we sought to identify known electrophysiological signatures (as established in rodents and primates and in some cases humans) from hippocampus and cortex of sheep either while they were performing cognitive tasks or during simple naturalistic behaviours. Using tried-and-tested neuronal recording techniques (individually adjustable tetrode electrode arrays) we were able to record local field potentials and multi neuronal activity for several months. Thus, we show that the technique is well suited for studies that require chronic recordings for longitudinal characterisations. As well as validating the methodology, we revealed for the first-time electrophysiological signatures of sheep hippocampal function that are largely consistent with the known properties for this thoroughly studied brain structure. As in other species, we observed a prominent theta rhythm oscillation which had also behavioural correlates. Unlike the case of rodents, however, where theta is mostly correlated with locomotion states, we observed that theta oscillations were associated with several other states, for example gaze fixation that has also been reported in non-human primates (Jutras et al., 2013). Furthermore, we observed nesting of gamma band oscillations by the slower theta oscillation phase that is seen in recordings of non-human primates. Gamma band activity also correlated with speed, as observed in rodents (Zheng et al., 2015).

Of particular note is that the frequency of the theta rhythm and sharp wave ripple oscillations are lower than that seen in rodents. This is consistent with a decrease in the dominant frequency of these oscillations as brain size increases (Buzsáki et al., 2013). While it is possible that the lower theta frequency is a result of the limited range of speed the sheep could attain within our experimental arena or that it is due to the fact that the electrode locations were in intermediate CA1 area and not dorsal CA1 (which is the hippocampal subfield that speed and oscillation correlations are observed in rodents (Schmidt et al., 2013), this latter explanation is inconsistent with the lower frequency or ripples, since ventrally in the rodent hippocampus ripples appear to be of faster frequency (Kouvaros and Papatheodoropoulos, 2017). Whatever the explanation, these two findings (lower theta and ripple oscillation frequencies) are of translational interest, given that low theta and ripple frequency rhythms are observed in hippocampal recordings from human subjects (Jacobs, 2014).

In the hippocampus, we observed putative neurons that were modulated by place but also a cell that exhibited rate remapping after manipulation on sensory cues. This extends the number of species in which the hippocampus has been implicated in spatial or episodic memory. The presence of place neurons in sheep hippocampus is not in itself surprising, but their existence opens up new vistas of discovery. Conducting studies on neurons that are modulated spatially are limited by the dimensions of the environment in which it is possible to test animals. This is particularly challenging when using non-human primates rather than rodents, since not only are they larger, but also their natural behaviour takes place in 3 dimensions. Sheep primarily move in 2 dimensions, so the technological challenges of capturing their behaviour are reduced. There are also a number of other advantages of using sheep for such studies. Because sheep are docile and easily confined by conventional farm fencing, they could potentially be studied, not only in large spaces but also in groups embedded in social contexts. Furthermore, sheep can easily perform complex cognitive tasks that measure decision making and the repertoire of behaviour that can be studied is extensive. Thus, an interesting opportunity arises for study mechanisms memory or spatial coding properties of the temporal lobe. These could be large scale, not only in terms of neural space (because of the size of the sheep brain), and in physical space (since sheep can be used outside the constraints of a normal laboratory environment) but also in terms of group size.

All recordings described here took place within the middle anteroposterior extent of the ectolateral gyrus with the largest number of putative cells being recorded from the third parietooccipital convolution overlaying the dorsal hippocampus. While a detailed functional anatomical atlas for the sheep brain does not yet exist, previous investigations associate the ectolateral gyrus with primary visual functions (Clarke and Whitteridge, 1976; Karamanlidis et al., 1979; Rose, 1942). In our investigations we observed highest decoding accuracy for behavioural state that was locomotion versus stationary state (during reward consumption within the two-choice discrimination task). We suggest therefore that this sheep brain area is involved in sensory-motor processing. Further investigation will be needed to clarify this.

Whilst our study serves as an example of the potential of large-scale electrophysiological recordings it also raises the potential for conducting imaging studies in animals embedded in naturalistic and social environments. The opportunity afforded by the ability of the sheep to carry substantial head-mounted devices combined with their lack of ability to interfere with such devices will permit studies that may be impossible to conduct in either primates or rodents.

Another area of research where large animals such as sheep are beginning to have impact is in testing of therapeutic interventions, such as gene therapies for neurodegenerative conditions (Evers et al., 2018; Morton, 2018; Pfister et al., 2018). Both the challenge of ‘scaling up’ to a larger brain and the availability of sourcing subjects can be met by using sheep, that are readily available in most countries.

In conclusion we demonstrated here the usefulness of a sheep as a large brain mammal for longitudinal electrophysiological studies at a single neuron resolution. The method serves only as an example, but if expanded with more modern electrophysiological tools the opportunity for large-scale monitoring of brain activity in a large unrestrained mammal is unprecedented. We suggest that although this species will not replace common laboratory species it could play a unique role in accelerating progress towards understanding normal and abnormal brain function.

## Acknowledgements

We thank Mr Roger Mason, and Dr Polly Taylor for anaesthesia and other technical support and Dr Matt Jones for guidance in establishing the recording technique.

---

**Supplementary Figure 1.**
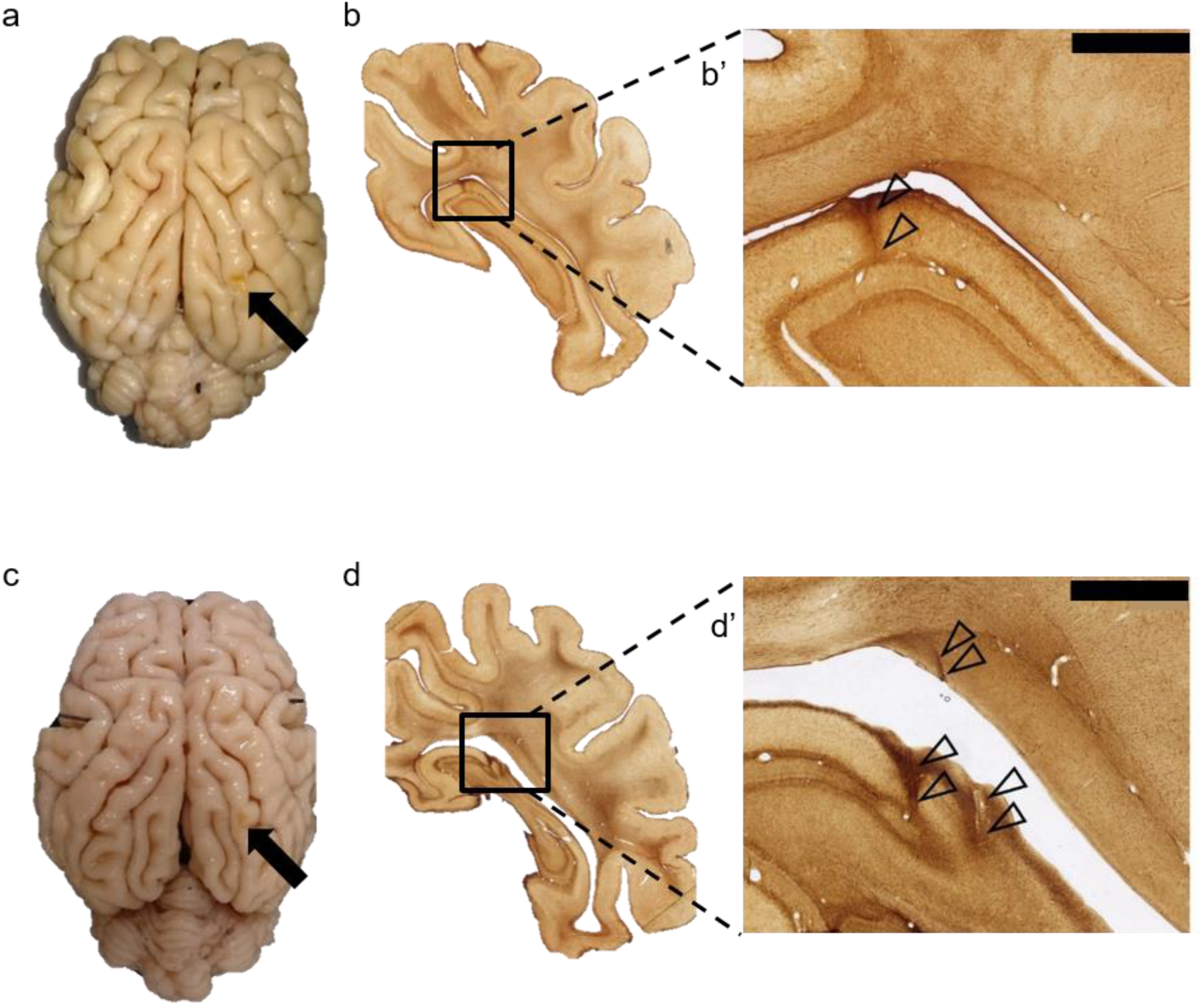
(a,c) Extracted brains from sheep 1 and sheep 2. Black arrows show the point of tetrode bundle penetration through the ectolateral gyrus. (b,d) Example histological evidence of electrode positions within the HPC formation in both sheep. Zoomed in sections (b’ and d’) with superimposed black triangles highlight electrode tracks. Panel (d’) is same as in Fig. 1. Scale bars in (b’,d’) correspond to 2.5mm.

**Supplementary Figure 2.**
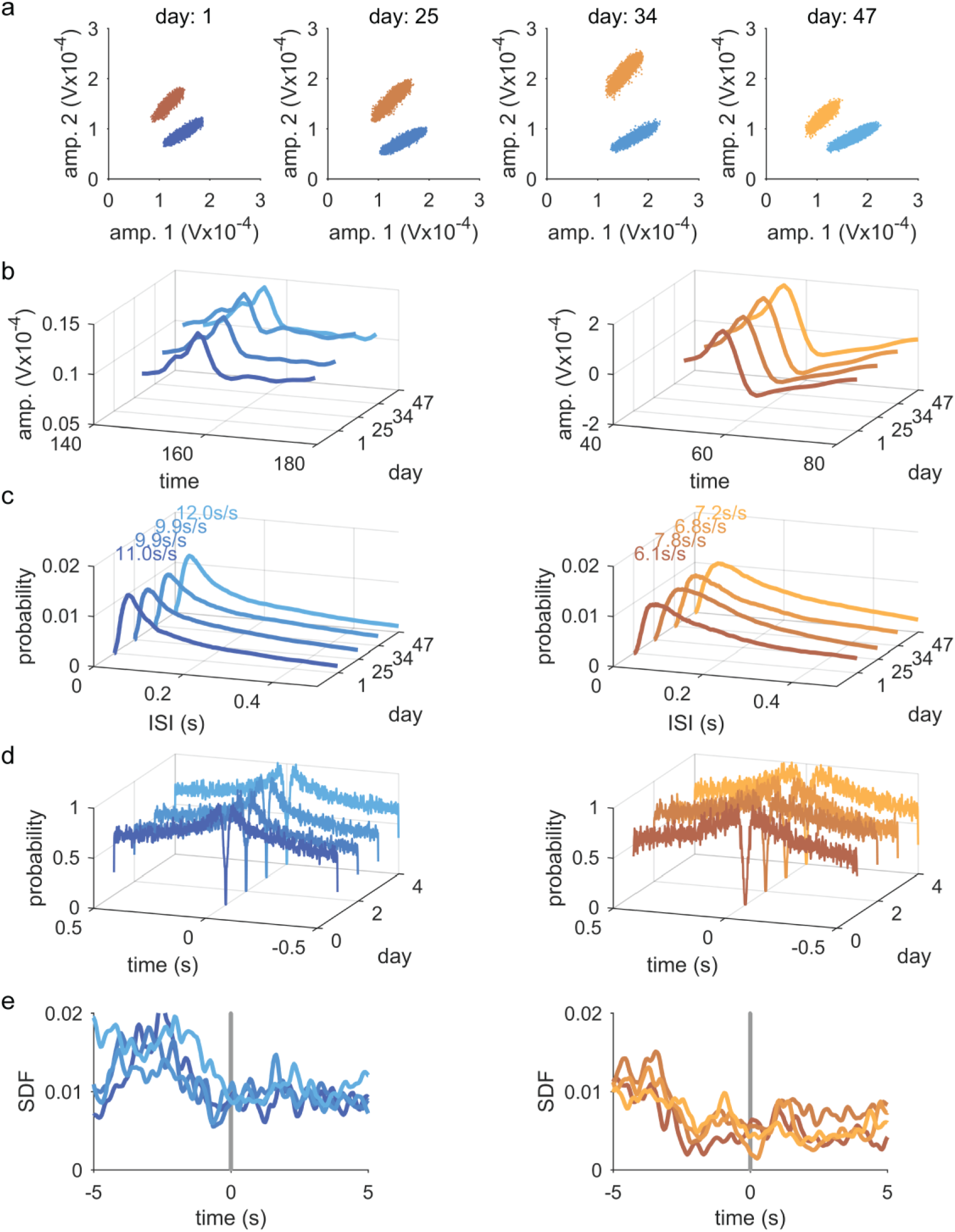
Example longitudinal recording from the same cells, extending across multiple weeks. Cluster features are constant across the different recording days. For each of the two cells identified on a single tetrode longitudinally across time (cell 1 blue, left panels; cell 2 orange, right panels): (a) amplitude clusters across two of the four tetrode wires (noise cluster suppressed for clarity). (b) mean spike waveforms on the first wire of the tetrode. (c) inter-spike interval plots. (d) normalised autocorrelograms. (e) spike density function (SDF) with a gaussian smoothing kernel of 50ms triggered onto the correct choice time points. Mean correlation coefficient of all combinations of cross-correlation functions (not shown) was 0.8 ±0.023 SEM, confidence interval, CI = 0.82 - 0.91, P = 8.9x10^-8^.

**Supplementary Figure 3.**
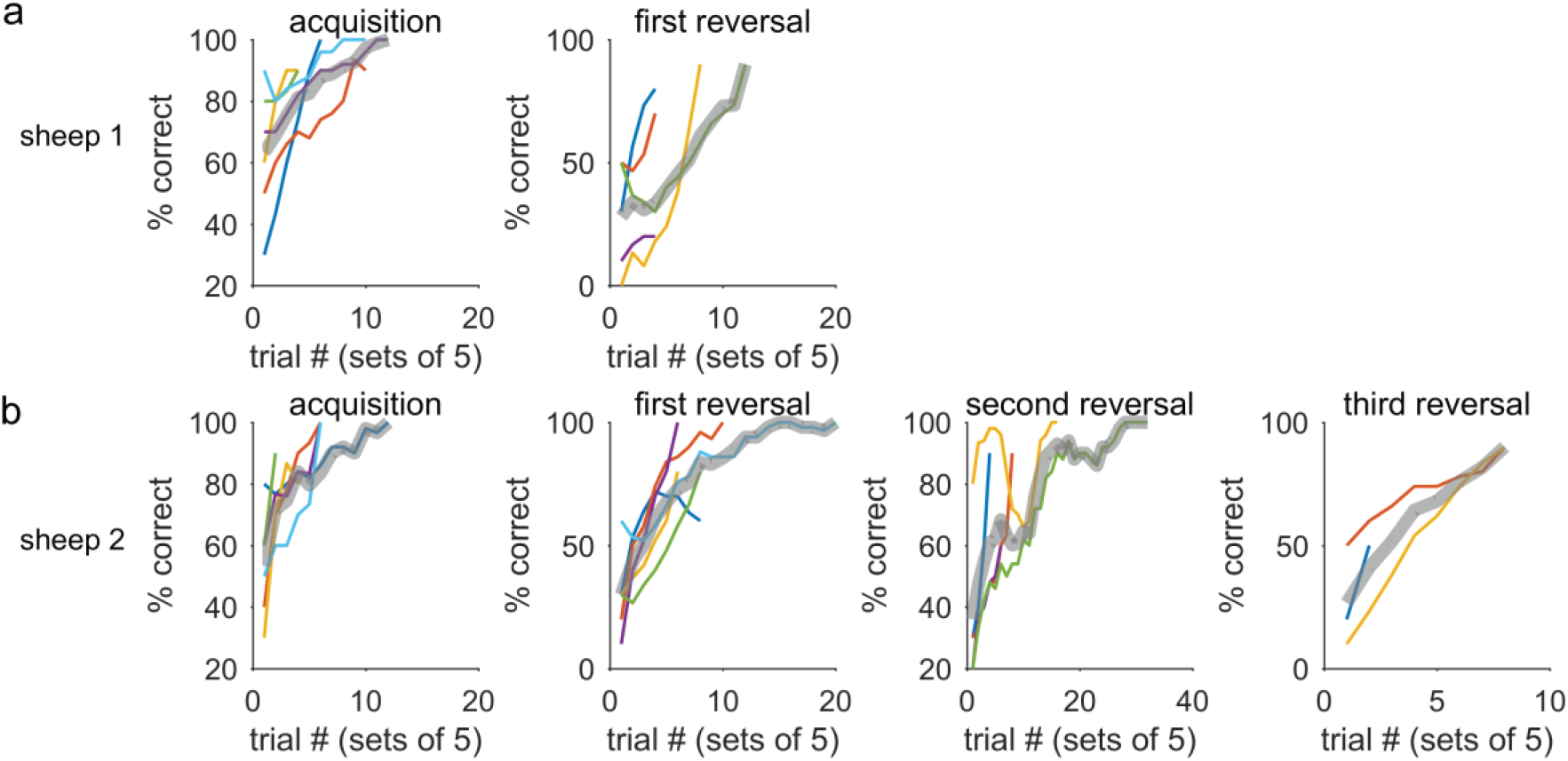
Example performance in two-choice discrimination acquisition and (serial) reversal for sheep 1 (a) and sheep 2 (b). Grey thick line in each subplot is the mean of several sessions with unique stimuli. Unique sessions are represented by the rest of the colors. Each datapoint represents the average of 5 consecutive trials. Data are smoothed with a moving average of window length 5. Reversal 2 and reversal 3 refers to identical symbols and repeatedly swapped contingencies.

**Supplementary Figure 4.**
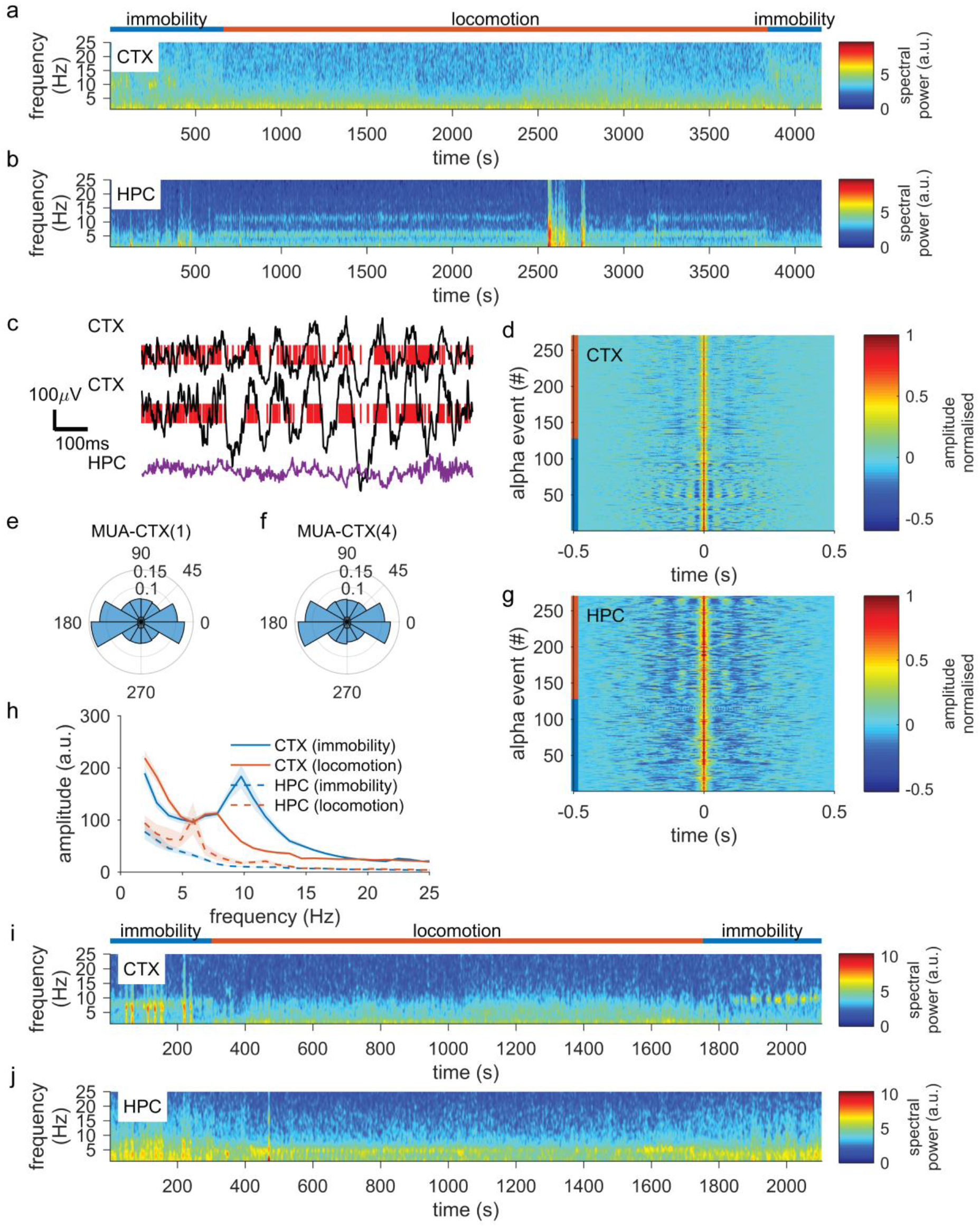
CTX alpha band oscillations are observed preferentially during immobility. Data in (a-e) are from sheep 1. (a) Spectrograms from CTX (top) and HPC (bottom) during immobility and cognitive testing. A 10Hz band is visible during immobility only. Theta state is suppressed during alpha episodes. (b) Example LFP from CTX and HPC during an alpha episode. Red ticks represent multiunit spiking activity. (c) All detected alpha events centered on the maximum amplitude point of each event (0s). Color scale represents normalised amplitude. (c) Polar histograms of multiunit spiking from two example tetrodes. Angle represents the phase of the alpha oscillation. (e) Average spectra from the two behavioural states for CTX and HPC. (f) Spectrograms equivalent to (a) but for sheep 2. The remaining equivalent panels are presented in the main text.

**Supplementary Figure 5.**
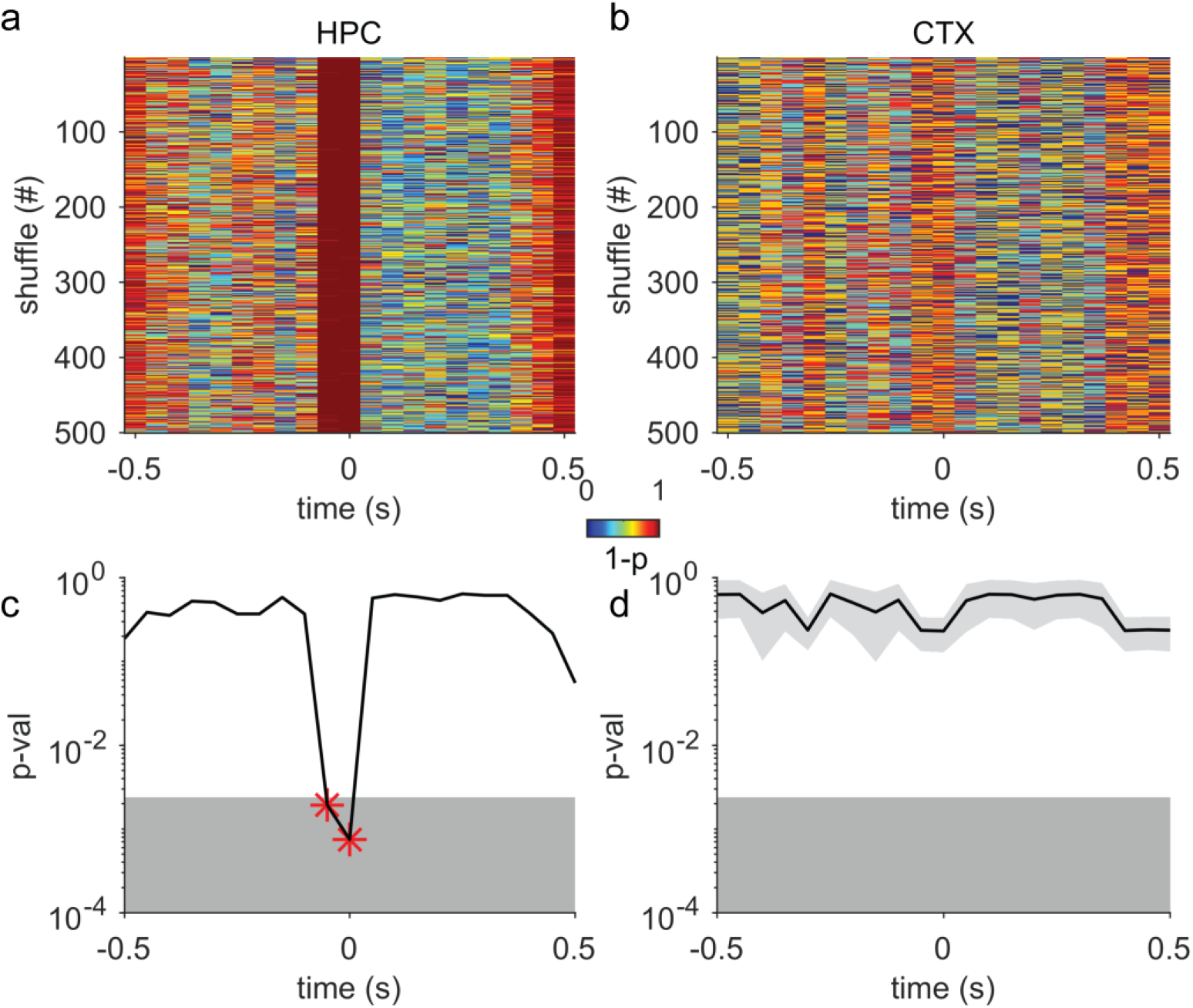
Shuffle test analysis for ripple triggered rasters in hippocampus and cortex. (a,c) Data are linearly shifted (independently and randomly for each trial) and then compared to the real data through a Wilcoxon signed rank test. Color represents significance value observed for each time bin and shuffle instance (colorscale is the p-value subtracted from unity so that red signifies smaller p values). (c,d) The distribution of shuffle tests reaches significance for the hippocampal activity and only around the ripple time point. In (c) red stars depict time bins where significance is reached. Time bin resolution is 50ms.

**Supplementary Figure 6.**
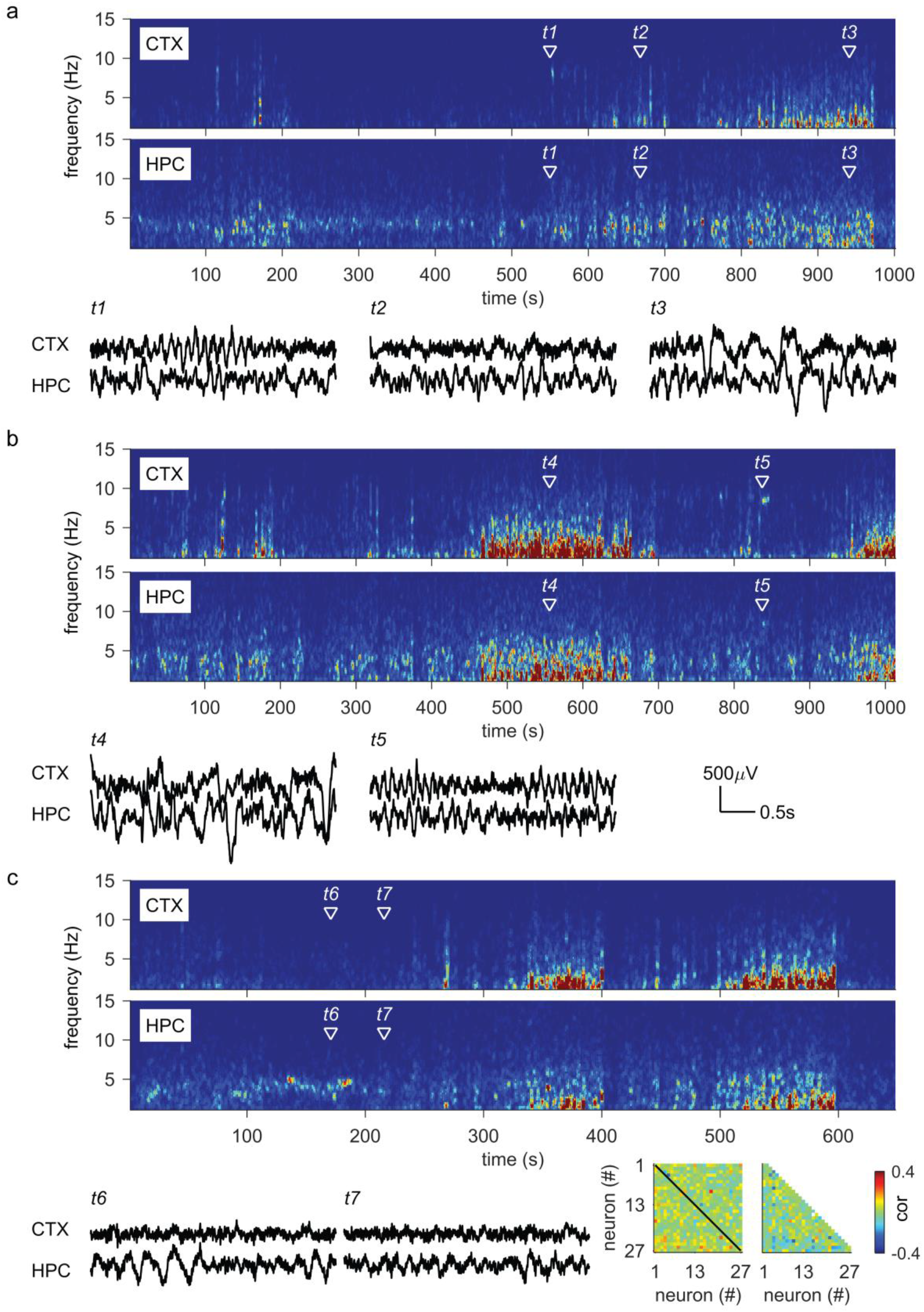
Examples from overnight recordings contrasting rapid eye movement and non-rapid eye movement sleep. (a-c) Examples from an overnight recording during (a) early night sleep episode, (b-c) late sleep episode. Spectrograms show the spectral content of electrodes in the putative locations of cortex (CTX; top spectrograms in a-c) and hippocampus (HPC; bottom spectrograms in a-c). Time points of interest are labelled as t1-t7 in both spectrograms and are also shown as time traces below each set of spectrograms. (t1: An alpha-band cortical oscillation while with the sheep seated with head erect, t2: A rumination episode with the sheep seated with head erect, t3: Early NREM episode with low amplitude slow-wave activity, t4: Large amplitude slow-wave sleep with head reclined and eyes closed, t5: Alpha band cortical oscillations following arousal and eyes opening while head is reclined, t6-t7: Putative REM sleep episode with theta oscillations in HPC and desynchronised CTX LFP while the sheep has head reclined is immobile and its eyes are closed. (d) Left color plot: Firing rate cross correlations between all pairs of recorded SUs for NREM sleep (lower triangle) and REM sleep (upper triangle). Black diagonal line delineates the upper and lower triangles. Right color plot: Difference in cross correlations between NREM and REM episodes. Color scale applies to both graphs. Calibration bars in (b) apply to all LFP waveforms (t1-t7). In (d) cross correlations are computed from intervals 330-420s and 510-610s for NREM and 123-257s for REM of the recording segment depicted in the spectrograms of (c).

**Supplementary Figure 7.**
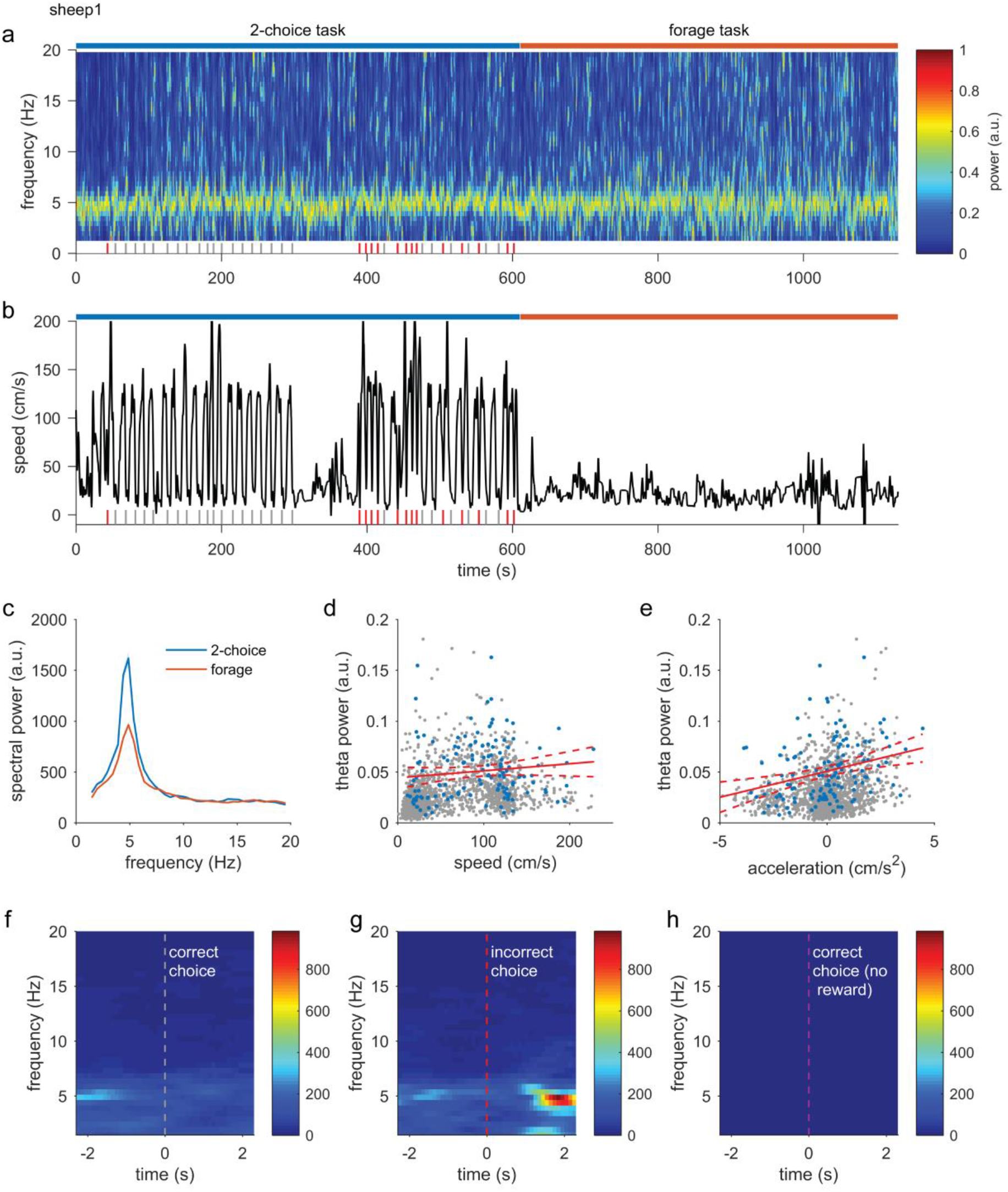
Example session from sheep 1 used for quantification of the hippocampal (HPC) theta power with speed of locomotion relationship. (a) HPC whole session spectrogram containing a period of two choice discrimination performance (blue bar above spectrogram) and a period of pellet foraging (red bar) with the latter typically having a low speed profile. (b) The speed of locomotion for the same period shown in (a). Ticks at the bottom of (a,b) show the timepoints at which the sheep triggered one of the choice sensors. (c) Average spectral power of HPC LFP for the two behavioural intervals. (d)Theta power - speed regression. Grey points: excluded intervals (delta-theta criterion ratio), blue points: included data, solid red line: linear fit, dashed red line: confidence intervals. (e) as in (d) but for acceleration. (f-h) triggered spectrograms around the choice timepoints. (f) correct trials, (g) incorrect trials and (h) correct but not reward trials.

**Supplementary Figure 8.**
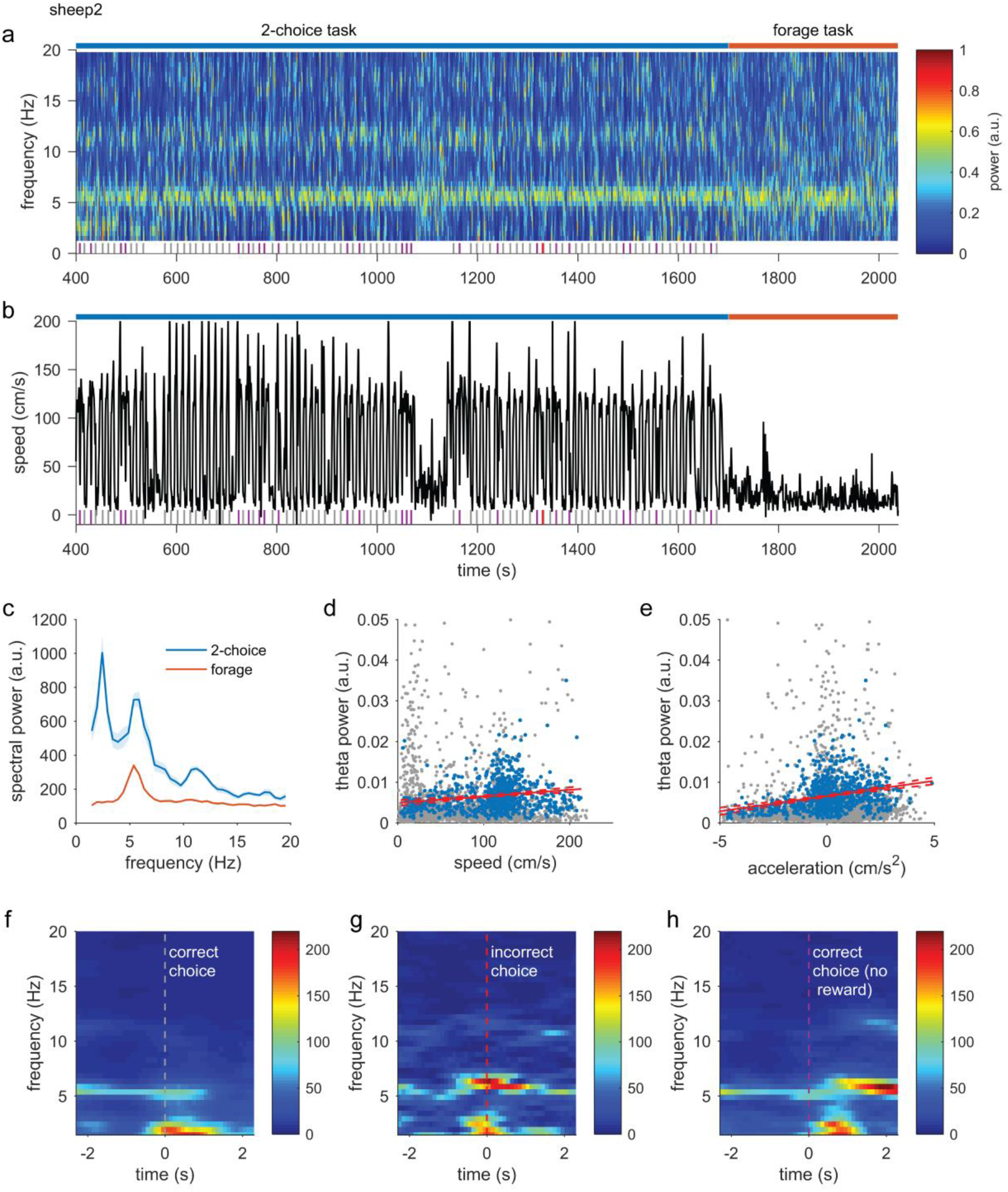
Example session from sheep 2 used for quantification of the hippocampal (HPC) theta power with speed of locomotion relationship. (a) HPC whole session spectrogram containing a period of two choice discrimination performance (blue bar above spectrogram) and a period of pellet foraging (red bar) with the latter typically having a low speed profile. (b) The speed of locomotion for the same period shown in (a). Ticks at the bottom of (a,b) show the timepoints at which the sheep triggered one of the choice sensors. (c) Average spectral power of HPC LFP for the two behavioural intervals. (d)Theta power - speed regression. Grey points: excluded intervals (delta-theta criterion ratio), blue points: included data, solid red line: linear fit, dashed red line: confidence intervals. (e) as in (d) but for acceleration. (f-h) triggered spectrograms around the choice timepoints. (f) correct trials, (g) incorrect trials and (h) correct but not reward trials.

**Supplementary Video 1.** Overhead view of the experimental arena during two-choice discrimination with concurrent electrophysiological recording. The sheep (sheep 2) is seen approaching one of two stimulus/reward locations sometimes after some deliberation. A high pitch auditory tone can be heard during incorrect choices (white triangle shape on black background) while another lower pitch sound is heard at the beginning of each new trial. The sound that immediately follows the correct choice (latin character L rotated by 90°) is that of the reward being dispensed below the local screen monitor.

**Supplementary Video 2.** A close up of sheep 2 during an approach to the correct stimulus screen. Tethering along with head implant, LEDs for tracking are clearly visible. Also visible is the optical sensor above the pellet dispenser box.

**Supplementary Video 3.** Sheep 1 is seen here performing a serial two-choice discrimination task during electrophysiological data acquisition. At the far end of the room are also visible two computer monitors that are displaying the data being acquired. Apart from the auditory tones being heard is also the sound of the spiking activity from a hippocampal tetrode. At 38s spiking activity from a putative place cell is heard and can also be seen on the right most computer screen.

**Supplementary Video 4.** Sheep 2 performing a simpler version of a two-choice discrimination task where it is required to approach the screen that displays a symbol versus the blank screen. Shown on the computer monitor is the local field potential trace with superimposed spiking events from a hippocampal tetrode. A clear and prominent oscillation (theta range) is seen to emerge upon the beginning of the locomotion episodes.

## Online Methods

### Animals

Two Welsh Mountain sheep (ewes aged 17 and 19 months at implantation) were used in this study. Both were obtained from a University of Cambridge stock flock. All procedures were conducted in accordance with the UK Animals Scientific Procedures Act (1986) and the University of Cambridge Animal Welfare and Ethical Review Bodies (AWERB).

### Surgery and animal handling

For several weeks prior to surgery, the sheep were handled routinely. They were trained to walk on a leash in order to habituate them to their human handlers and to be comfortable when isolated from the flock. Both sheep were content to work in isolation from conspecifics in the presences of their handler(s).

Sheep were food-deprived overnight prior to surgery. Anaesthesia was induced using intravenous administration of Alfaxalone (Alfaxan, Jurox, UK) at 3mg/kg and maintained using 2-3% isoflurane through a Manley ventilator. Throughout surgery, end-tidal CO2 and mean arterial blood pressure were kept between 25-30 mmHg and 70-90 mmHg, respectively. Vital functions were recorded at regular time intervals throughout the procedure. Intravenous fluids were supplied at a rate of 5 ml kg^-1^hr^-1^ (lactated Ringers, Hartmann’s Solution 11 by Aquapharm). Using aseptic techniques, the dorsal surface of the skull was exposed and cleared of any connective tissue using a bone scraper. The skull was cleaned with 3% hydrogen peroxide, rinsed with sterile saline and any remaining connective tissue was then removed. A 15 mm by 15 mm craniotomy was drilled on the right hemisphere at a distance of 9 mm posterior and 8 mm lateral from the intersection of the frontal and posterior skull bones (Figure X). Once the craniotomy was completed, the dura mater was incised using a dura incision tool at a carefully selected location to avoid any vessels. With visual access to the cortex, the implant site was selected at approximately the anterior third of the ectolateralis gyrus at a location with little to no vasculature. As the pia matter was also found to impede tetrode penetration, it also was incised. This was performed under a microscope using electrocautery micro-forceps. In case of bleeding, Gelfoam was applied briefly into the craniotomy. Using a stereotaxic manipulator, the micro-drive implant was lowered in place, with the bottom of the large cannula at close proximity to the surface of the brain (less than 1 mm distance). A ground wire was then connected to the skull through a stainless-steel screw. At this time point, data acquisition was enabled, and electrodes were driven by 1-2 mm into the first layers of the cortex. This was deemed necessary to ensure that electrodes were able to move freely prior to fixating the implant onto the skull permanently. Once penetration was confirmed, the craniotomy was padded with an absorbable hemostat (SURGICEL® FIBRILLAR™, Ethicon, Johnson & Johnson), dried off of any cerebrospinal fluid and then sealed with gentamicin containing dental acrylic (Depuy, Johnson & Johnson, p/n 3325020, CMW1). The acrylic was allowed to increase in viscosity before application in order to avoid the possibility of leakage through the absorbable hemostat layer. With the tetrode microdrive fixed in place on the skull, the electrode drive chamber was attached to the cranium using stainless steel screws and then covered with dental acrylic. The wound was finally sutured using the horizontal and purse string suture techniques. Finally, the wound was sealed using wound plaster spray (Kruuse Wound Plast, Denmark). With all invasive procedures completed, the sheep’s head was detached from the stereotaxic frame, and isoflurane, nitrous oxide and intermittent positive pressure ventilation were discontinued. Oxygen flow was maintained while respiration was supported manually until spontaneous breathing started. The sheep was then placed in a recovery area with padded walls and once the anesthetist deemed appropriate, the intubation tube was removed. Once the sheep was alert enough to stand, it given pelleted feed and then was returned to its home environment. The sheep was monitored regularly for the rest of the day. A four-day course of antibiotic treatment (penicillin, Depocillin, 15mg/kg), non-steroidal anti-inflammatory treatment (carprofen, Rimadyl, 4mg/kg, once per day for 2-3 days) and pain relief treatment (Buprenophrine, Vetergesic, 0.01mg/kg, once per day for 2-3 days) was given to each sheep post-operatively. In addition, a single dose of dexamethasone was given within the first 24 hours after surgery (Dexadreson; MSD Animal Health, UK; 2mk/kg). More details for animal habituation and surgery techniques can be found elsewhere(Perentos et al., 2016b).

### Tetrode microdrives and electrode chambers

We used custom-built tetrode microdrives, similarly to those described by Nguyen et al.(Nguyen et al., 2009). The main difference was that the travel distance was set to 2 cm thus requiring longer screws and cannulae to accommodate for the extra travel. Tetrodes were manually assembled from 12.5µm diameter Nichrome wire, coated with ¼ Hard PAC (Kanthal Precision Technology, Palm Coast USA). Once electrodes were loaded into the Microdrive, they were attached to Neuralynx interface boards. Two days prior to surgery, electrodes were gold plated to achieve a target impedance of 250kΩ, suitable for spike detection. To protect the tetrode microdrive against physical force an electrode chamber was used. It was a requirement that the chamber could be easily installed after the tetrode microdrive was cemented in place. Therefore, the chamber consisted of three parts, a lid and a two-part snap-on conical shaped chamber (Fig. 1b). The chamber was 3D printed out of polyamide (Quality Equipment, G. E. Baker, UK Ltd) and attached to the skull through several stainless-steel flanges.

### Electrode movement

Four (sheep 1) and seven (sheep 2) days after surgery, electrode movement commenced. The sheep was restrained using a Gambrel restrainer (Wynnstay Agriculture, UK), and encouraged to stay still using intermittent reinforcement (food pellets). Electrodes were then advanced by 150-300 µm each per day. After 7-10 days post-surgery, daily recordings commenced. At the end of each recording session the tetrodes were advanced, until they eventually reached the hippocampus. Local field potential and single unit sampling continued from tetrodes left in the superficial layers of the cortex as well as those that had been directed down to the hippocampal formation.

### Recordings and equipment

All recordings took place in a large dark farm shed containing a specially constructed spatial arena (Fig. 1f,g) that was fitted with an automated end-to-end two-choice discrimination apparatus. This apparatus was controlled via a Matlab (The MathWorks, Inc., Natick, MA, USA) and a digital acquisition device (USB-1208fs, Measurement Computing Corporation). It allowed two-choice discrimination trials at either end of the arena in an alternating manner, thus forcing the sheep to traverse the environment for reward opportunities. A variant of this behavioural apparatus that implements a single two-choice discrimination is described by McBride et al.(McBride et al., 2016). Briefly, the apparatus consists of infrared beam sensors, computer monitors, automatically triggered feeders and a digital acquisition device, all controlled through scripts written in Matlab. Electrophysiological recordings were obtained using the Digital Lynx SX system (Neuralynx Inc. Bozeman Montana). All tetrode channels were sampled at 32 kHz while one electrode from each tetrode was used to obtain local field potential recordings at a sampling rate of 8 kHz. Customised battery driven LED lights were attached to the sheep’s head and were used for tracking purposes. Four overhead cameras (Jai CV-3200 NTSC with a Fujifilm DF6HA-1B 6mm lens) were used to track the location of the sheep in the arena. Camera sampling rate was 25 fps and the Neuralynx VTS software was used to track the sheep’s position. The physiological and behavioural acquisition computers were synchronised through the external TTL ports of the Digital Lynx SX system.

### Behavioural testing

In most recording sessions the sheep performed several tasks. They performed a simple stimulus association task, whereby they were required to approach the computer monitor that displayed a symbol in order to attain a reward. They also performed two-choice discrimination tasks, sometimes followed by reversal of contingencies. In some trials, sheep engaged in a spatial exploration task, where sheep would search for food pellets (used as a reward) that were scattered on the floor of the arena. In each recording session, we recorded resting-but-awake phase activity at the beginning and at the end of each recording session since the sheep was invariantly restrained for attachment/detachment of cables and preamplifiers. During these recording phase we searched for correlates of inactivity such as sharp wave ripple oscillations in the hippocampus, or neocortical alpha band oscillations. Whole night electrophysiological recordings were also obtained from each sheep.

### Typical recording session process

In a typical session, a sheep was transferred to the shed containing the recording area on a lead or by voluntarily following the experimenter (Video 1). Inside the shed, but outside the arena the sheep was restrained using a Gambrel restrainer (Wynnstay Agriculture, UK). The electrodes were connected to the recording equipment via the head mounted preamplifiers and a protective dome was attached to house the preamplifier assembly. LED lights were attached to the dorsal aspect of the head and a 5-minute baseline recording was obtained. The sheep would then be guided into the experimental arena, and the behavioural testing would commence. Typically, the sheep would perform 80-100 two-choice discrimination trials that sometimes contained reversal of contingencies. Finally, pellets would be scattered on the floor of the arena in order to encourage the sheep to visit areas of the arena that it would not otherwise visit during the two-choice discrimination task.

### Histology

At the end of the experiments the sheep were euthanized, and their brains examined to establish the location of the electrodes. Sheep were given a lethal dose of pentabarbitone and once death was confirmed the jugular veins were catheterised and flushed with 2L of saline. The brains were then fixed with 4% paraformaldehyde (Sigma). The brain was then extracted from the skull and immersed in 4% paraformaldehyde for a month. After that, it was then transferred into a cryopreservation solution (20% sucrose). All membranes covering the cortex and cerebellum were removed and the brain was cut into the two hemispheres and further blocked around the area of tetrode penetration thus facilitating faster sucrose penetration. The tissue was then cut into 60µm sections and alternate sections were stained with cresyl violet or immunostained for glial fibrilliary acidic protein (using 1:1000 anti-GFP antibody, VectorLabs). The former was used to visualise electrode tracks and the latter was used to examine for the presence of reactive astrocytosis (Suppl. Fig. 1).

### Data analysis

Data analysis was performed in Matlab, Mathworks®, using custom made scripts as well as other toolboxes such as Chronux. Video trackers were stitched together manually thus combining the four video tracking feeds into a single continuous trace. Spike sorting was performed using MClust v4.4. Clusters were curated manually using available features such as amplitudes and principle components. Clusters with contaminated refractory periods were not considered for further analysis. For decoding purposes 8-second worth of firing rates (binned at 100ms) centered on choice points from all cortical cells were used. Decoders were implemented on a per session basis thus avoiding collapsing data from different sessions. Linear discriminant analysis was implemented using the ‘fitcdiscr’ built-in Matlab function with parameters of diagonal covariance matrix and a 10-fold cross-validation. Cross-validation error (kfoldLoss matlab function) was used as a metric to compare the models of each of the predictors. Predictors were also averaged within the same 100 ms windows. Throughout, unless otherwise stated, mean data are shown as mean ± SEM.

For ripple detection, all tetrodes identified as putatively hippocampal were used. LFP traces were high pass filtered at 100 Hz and z-scored. The smoothed and rectified Hilbert envelope was then used for detection of large amplitude events (z-score < 3). To look for the concurrent presence of a sharp wave event coincident with above detected events raw LFP data were used after low pass filtering as 15 Hz. To assess the frequency content of these events triggered spectrograms centered on the detected peaks were computed using the mtspectramc Chronux function with taper parameters set as 3 and 5 and a window length of 150ms.

For the detection of alpha band oscillations during periods of inactivity, a subset of putatively cortical tetrodes from the two sheep were used. Threshold crossings (z = 3) from narrowband (6-12 Hz) and z-scored LFPs were detected, a local search was performed at either side of these detection to establish the extents of the detected events. Overlapping events were joined together. Detections were quantified as a function of behavioural state (behaviioral task in the arena versus restraint epochs outside the arena). Similarly, slow-wave oscillations were detected using low z scored and pass filtered (< 3 Hz) cortical LFP data (and HPC for comparison purposes). Threshold was set at z = 5 SDs. In order to assess the frequency content accompanying slow wave events, multitaper triggered spectrograms around the negative peaks were computed (window size =1.5s, tapers (1,3)). Assessments of relationships between speed and theta or gamma oscillations in the HPC where conducted using linear regressions using the fitlm matlab function. Cross-frequency coupling was assessed using the modulation index method (Tort et al., 2010) and shuffle correction was achieved through phase randomisation of the narrowband filtered LFPs (1000 surrogate datasets whose mean comodulogram was subtracted from the real comodulogram before further speed group comparisons). Spike phase modulation by an underlying oscillation was derived through interpolation between successive peaks and troughs detected from the narrow band LFP signal. Spatial modulation of hippocampal cells was assessed as follows: First spatial occupancy was computed with a resolution of 25cm^2^ as well as the total number of spikes within each corresponding spatial bin. Spike counts were then normalised by the occupancy and filtered with a two-dimensional gaussian smoothing kernel with standard deviation of 0.3. Bins with low occupancy values (< 0.3s) were excluded from the analysis. The information content metric was used to quantify the spatial selectivity of identified cells (Skaggs et al., 1993).

